# Chronic variable mild stress alters the transcriptome and signaling properties of the anterodorsal bed nuceleus of the stria terminalis in a sex-dependent manner

**DOI:** 10.1101/2024.11.11.623087

**Authors:** Thomas J Degroat, Sarah Paladino, Benjamin A Samuels, Troy A Roepke

**Author notes:** **Corresponding Author:** TA Roepke Permanent address: Department of Animal Sciences, School of Environmental & Biological Sciences, Rutgers, the State University of New Jersey, 84 Lipman Drive, Bartlett Hall, New Brunswick, NJ 08901.

## Abstract

Chronic stress is a physiological state marked by dysregulation of the hypo-pituitary-adrenal axis and high circulating levels of stress hormones, such as corticosterone in mice or cortisol in humans. This dysregulated state may result in the development of mood disorders but the process by which this occurs is still unknown. The bed nucleus of the stria terminalis (BNST) serves as an integration center for stress signaling and is therefore likely an important area for the development of mood disorders. This project utilized a chronic variable mild stress (CVMS) paradigm to persistently stress mice for 6 weeks followed by RNA-Sequencing of the anterodorsal (ad) BNST and electrophysiology of corticotropin releasing hormone-expressing cells in the adBNST. Our results show significant sex-biases in the transcriptome of the adBNST as well as effects of CVMS on the transcriptome of the adBNST specifically in males. Female biased genes are related to synaptic transmission while male biased genes are related to RNA processing. Stress sensitive genes in males are related to synaptic transmission and synapse formation. Additionally, electrophysiology data showed that CVMS suppressed the M-current in males but not females. However, CVMS increased the strength of excitatory post-synaptic currents in females but not males. This suggests significant differences in how males and females process chronic stress. It also suggests that the BNST is more sensitive to chronic stress in males than in females.

## Introduction

In the central stress response, the major signaling molecules include corticotropin releasing factor/hormone (CRF/H) and adrenergic neurotransmitters. By contrast, the major stress molecule in the periphery is cortisol in humans and corticosterone in rodent models. When presented with a stressor, the brain must first recognize the stressor through the integration of multiple processing centers, notably the amygdala, paraventricular hypothalamus, and locus coeruleus (Herman et al., 2005; Poe et al., 2020). The amygdala and paraventricular hypothalamus produce CRF/H, while the locus coeruleus produces norepinephrine and responds to CRF/H. CRH from the paraventricular hypothalamus controls the hypothalamic-pituitary-adrenal axis.

The bed nucleus of the stria terminalis (BNST), a part of the limbic system or extended amygdala, is a region that also produces CRF-mediated output to mood regulation. Many of the previously discussed stress-related regions direct innervate the BNST and as such, it also can serve as an integration center for the central stress response (Herman et al., 2005). This heterogeneous area expressed receptors for various stress-related proteins, including CRF receptors and adrenergic receptors (Becker et al., 2008). With its relation to stress signaling, the BNST is implicated in the development of mood disorders. The BNST is of particular interest for studying the onset of mood disorders as previous studies suggest the BNST is only activated through long-context stressors, indicating that the BNST may specifically process chronic stress, a well-known factor that can precipitate mood disorders (Hammack et al., 2015).

When studying how chronic stress develops into a mood disorder, one must also consider how variability across the sexes influences responses to chronic stress. In clinical research, it has been shown that most mood disorders have a strong gender bias, with major depressive disorder being twice as prevalent in cisgender women than cisgender men (Bromet et al., 2011). Additionally, it is also known that the differing hormone profiles of cisgender men and women interact with depression differently as estrogen levels can be a significant regulator of depressive symptoms (Lai, 2011; Albert & Newhouse, 2019). In menstruating people, it has been shown that high estrogenic states, like the follicular phase of the menstrual cycle, can significantly worsen the severity of depressive symptoms i.e., in women with pre-menstrual dysphoric disorder (Freeman, 2003). However, estrogen is also important for mood regulation as low estrogenic states, like menopause or postpartum, can also lead to erratic moods or depression (O’Hara & McCabe, 2013; Li et al., 2024). Thus, it is important to understand how chronic stress is influenced by gonadal steroids and how gonadal steroids are influenced by chronic stress. In our previous publication, we found that the response to a chronic variable mild stress (CVMS) paradigm elicit different responses in avoidance behavior across the sexes and across the estrous cycle (Degroat et al., 2024).

To gain a better understanding of mood disorders, multiple levels of analysis are needed. Previously, we found that the CVMS paradigm also altered the M-current and excitatory post-synaptic currents (EPSCs) in BNST CRH neurons from male mice (Hu et al., 2020) but not in BNST NPY-expressing cells in either sex (Degroat et al., 2024). Further, when using bulk RNA-Sequencing of the anterodorsal (ad) BNST, we found no effects of CVMS on the transcriptome 18 h after the last stressor (Degroat et al., 2024). In the current study, we examined the influence of CVMS on CRH excitability, M-current activity, and EPSCs, in both male and female mice, and, due to the transient nature of the transcriptome, on the transcriptome of the adBNST 1 h after the last stressor (wet bedding). We postulate that the earlier collection post-stress would provide better insight into how stressors alter the transcriptome of the adBNST in a sex-specific matter that may lead to the sustained differences in neurophysiology. Our main goal is to characterize how the neurons of the adBNST, especially CRH neurons, respond to chronic stressors across the sexes.

## Materials & Methods

### Animals

All procedures were in accordance with National Institutes of Health standards and approved by the Rutgers Institutional Animal Care and Use Committee and agree with ARRIVE’s guidelines for reporting animal research. Mice used for all cohorts were adult male and female C57BL/6J mice. Mice used for electrophysiology were bred in house, mice used for RNA-Sequencing were purchased at 3 weeks of age from Jackson Laboratories (Cat# 000664). For electrophysiology, mice were cross-bred between an Ai14 parent and a Corticotropin Releasing Hormone (CRH)-Cre parent so that offspring had mutations for both Ai14 and CRH-Cre, resulting in exclusive expression of td-Tomato in CRH-expressing cells. Both Ai14 and CRH-Cre breeding mice were purchased from Jackson Laboratories (Cat # 007914 & 012704, respectively). Control or stress conditions were started at 7-10 weeks of age. Except for stressors, mice were housed in a temperature and humidity-controlled room (22°C, 30-70% humidity), on a 12/12 h light/dark cycle, and provided food and water *ad libitum*. Sample sizes of n=7-9 cells for electrophysiology allowed for detection of affects with a power = 0.8 at α = 0.05. This was based on previously published studies in our lab (Hu et al., 2016; Hu et al., 2020; Degroat et al., 2024). RNA-Sequencing was conducted via pooling 3 biological replicates into a single sample with 15 mice per group, resulting in an ultimate sample of n=5 per group. For electrophysiology mice, they were sacrificed the morning after the last stressor, approximately 18 hours, at 9:00am-10:00am. For the RNA-Sequencing cohorts, mice were euthanized 1 hour after the last morning stressor at 10:30am-11:30am.

### Chronic Variable Mild Stress Paradigm

The Chronic Variable Mild Stress (CVMS) paradigm is 6-week schedule of 1-2 minor stressors every day, typically with one morning stressor and one evening stressor. This paradigm follows the same procedure as outlined in our previous publication (Degroat et al., 2024). Briefly, mice were subjected to the following stressors: 3 or 8 hours of humid bedding, 8 hours of no bedding, 8 hours of replacing bedding with room temperature water, placing the cage on a 45° tilt for 8 hours, isolation for 8 hours or overnight, exposure to novel mice’s dirty cages for 8 hours, lights on overnight, 2 or 3 hours of dark during typical light cycle, 5 or 6 bedding changes in a single day, 2 minute swim stress in 4°C water, 4 minute swim stress in room temperature water, 15 minute cold stress, and 15 minutes of predator sounds, and 1 hour of restraint stress.

### RNA-Sequencing

To collect tissue for RNA-Sequencing, mice were first anaesthetized via ketamine followed by decapitation. Brains were dissected out, sliced, and then microdissected to isolate the adBNST. Tissue samples from 3 biological replicates were then pooled prior to RNA extraction and were stored in RNAlater at −80°C. RNA was extracted using an RNAqueous Micro Isolation kit (Invitrogen; AM1931). Integrity, concentration, and quality of the RNA samples were analyzed with an Agilent 2100 Bioanalyzer prior to being sent to the JP Sulzberger Columbia Genome Center (New York, NY) for sequencing. Any sample with an RNA integrity <8 was discarded. Libraries were constructed using Illumina TruSeq Stranded mRNA Library Prep Kit with KAPA HiFi HotStart Ready Mix for the final PCR step and were sequenced on an Element AVITI with 100bp paired-end reads at a depth of 40 million. Resulting fastq files were then analyzed as previously described (Degroat et al., 2024). RNA-Sequencing had a sample size of n=5 for each group with no outliers. Raw sequencing data can be accessed at GEO GSE281507.

### Electrophysiology

Electrophysiology was conducted as previously described (Degroat et al., 2024). Briefly, mice were decapitated between 8:30am and 10:00am. The brain was then dissected out and sliced on a VT1000 S vibrating blade microtome (Leica) at 250 nm thickness in a high sucrose artificial cerebral spinal fluid (aCSF) that was prepared in-house (containing in mmol: 208 sucrose, 2 KCl, 26 NaHCO3, 10 glucose, 1.25 NaH2PO4, 2 MgSO4 1 MgCl2, 10 HEPES with a pH of 7.35 and osmolarity of 290-310 mOsm). Slice were then allowed to acclimate at room temperature in a bath of continuously oxygenated aCSF (containing in mmol: 124 NaCl, 5 KCl, 2.6 NaH2PO4, 2 MgCl2, 2 CaCl2, 26 NaHCO3, 10 glucose with a pH of 7.35 and osmolarity of 290-310). Once acclimated, individual slices were placed in the bath of the electrophysiology rig that contained the same aCSF recipe as the acclimatization chamber. Individual cells were then selected by fluorescence of the tdTomato. A borosilicate pipette backfilled with an internal solution (containing in mmol: 10 NaCl, 128 K-gluconate, 1 MgCl2, 10 HEPES, 1 ATP, 1.1 EGTA, and 0.25 GTP with a 7.35 pH and 290-300 mOsm) and used to patch onto the cell. Membrane values and standard protocol was recorded for every cell and throughout the duration of the patching to ensure the health of the cell. Data collection was done using 1550B Axon Instruments Digidata acquisition system, 700B Axon Instruments Multiclamp amplifier, and pCLAMP software (version 11.1) from Molecular Devices. Analysis of the M-current was done on Clampfit (part of the pCLAMP 11 software suite) as previously described (Degroat et al., 2024). Excitatory post-synaptic current (EPSC) was analyzed on Easy Electrophysiology. EPSCs were detected using the template event detection feature using the deconvolution method with a theta of 4.5. Events were then individually checked to ensure noise was not accidentally counted. Events were then averaged, and number of events, average amplitude, and average area under the curve (AUC) were recorded. M-current tests had sample sizes of n=9 for control males, n=10 for stressed males, n=8 for control females, and n=9 for stressed females. There were no outliers for M-current. EPSC tests had sample sizes of n=7 for control males, n=8 for stressed males, n=8 for control females, and n=9 for stressed females. A table of outliers for EPSCs can be found as Supplementary Table 1. Females were tested only in diestrus to ensure consistent hormone profiles between mice,

### Statistics

Unless otherwise specified, all tests were analyzed using Prism (GraphPad Prism, Version 10, Dotmatics) and used a p-value or adjusted p-value of <0.05 for significance. Body weight tracking and electrophysiology tests were first checked for outliers by the Grubbs’ test using a critical value of 0.05 as the cut off. EPSC data were analyzed by a two-way ANOVA. RNA-Sequencing data was analyzed by DESeq2 on R Studio. Cumulative body weight gain and the IV plots of the M-current recordings were analyzed by a repeated-measures two-way ANOVA. Max peak of the current at −35mV, resting membrane potential, and change in input resistance were analyzed by a two-way ANOVA followed.

## Results

### Body Weight

Body weight gain was impacted by CVMS exposure in both the males and females; however, the sexes responded differently. In males, body weights between control and stressed counterparts stayed consistent for the first three weeks, then started to diverge (Fig. 1a). After week one, the control males gained 2.6% of their body weight and stressed males gained 3.3%. After week 2, control males gained 7.9% while stressed males gained 7.6%. After week three, control males gained 9.3% while stressed males gained 7.9%. After week four, control males gained 13.8% while stressed males gained 9.8%, (trending p= 0.054). After week five, control males gained 15.5% while stressed males gained 11.0% (p<0.0001). After the final week of stressors, control males gained 19.0% while stressed males gained 12.4% (p<0.0001). This data indicates that CVMS induces an inhibition in weight gain, presumably through changes in feeding or metabolism, in male mice.

**Figure 1:**
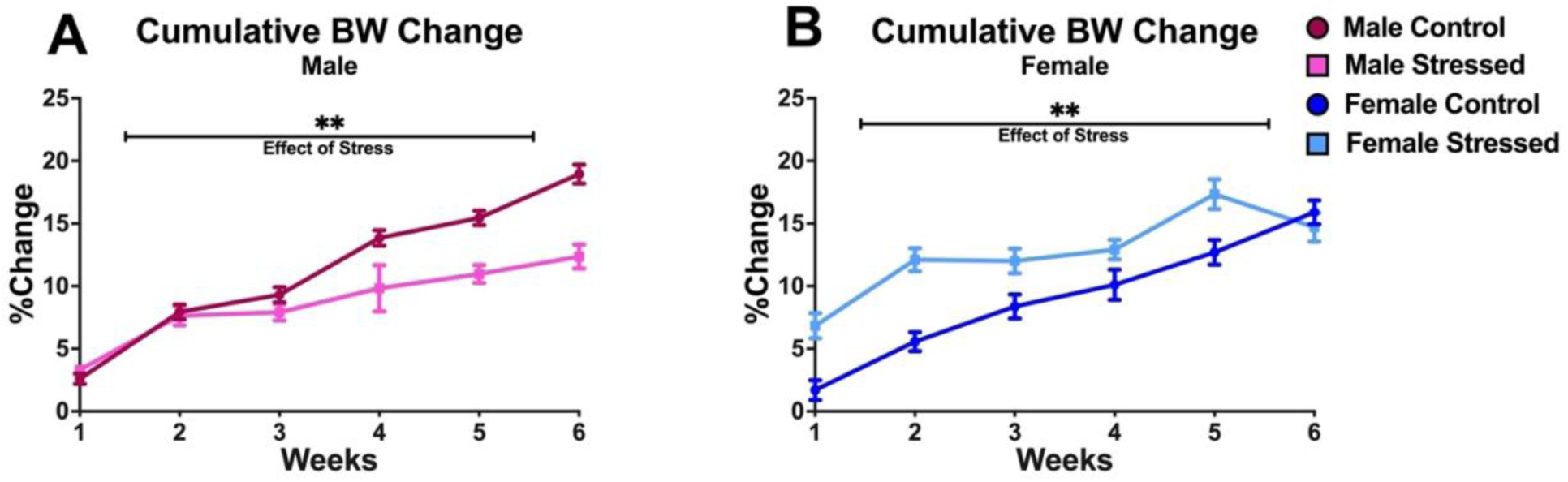
Cumulative %change in body weight during the 6-week stress paradigm in (A) male and (B) female mice. Analyzed by repeated measures two-way ANOVA. (*=0.05-0.01, **=0.01-0.001, ***=0.001-0.0001).

In females, body weight gain was different at the beginning, then converged towards the end of the paradigm (Fig. 1b). After week one, control females gained 1.7% of their body weight stressed females gained 6.8% (p=0.0004). After week two, control females gained 5.6% while stressed females gained 12.1% (p<0.0001). After week three, control females gained 8.4% while stressed females gained 12.0% (p=0.0135). After week four, control females gained 10.1% while stressed females gained 12.9% (not significant (ns)). After week five, control females gained 12.7% while stressed females gained 17.3% (p=0.0057). After the final week of stressors, control females gained 15.9% while stressed females gained 14.7% (ns). For females, data shows that they have a transient increase in body weight gain after exposure to stress, which plateaus under chronic conditions.

### adBNST Transcriptome

The principal component analysis of all samples used for RNA-Sequencing illustrates that there are no outliers for any groups as all samples fit within their respective 95% confidence interval ovals (Fig. 2a). It also finds that males and females are separating strongly, and male controls are separating from male stressed mice; however, the control and stressed female samples are almost entirely overlapping.There were 248 genes found to be significantly differentially expressed between males and females and a further 216 that were trending (0.10>p>0.05) (Fig. 2b). Of the significant genes, 116 were male biased while 132 were female biased.

**Figure 2:**
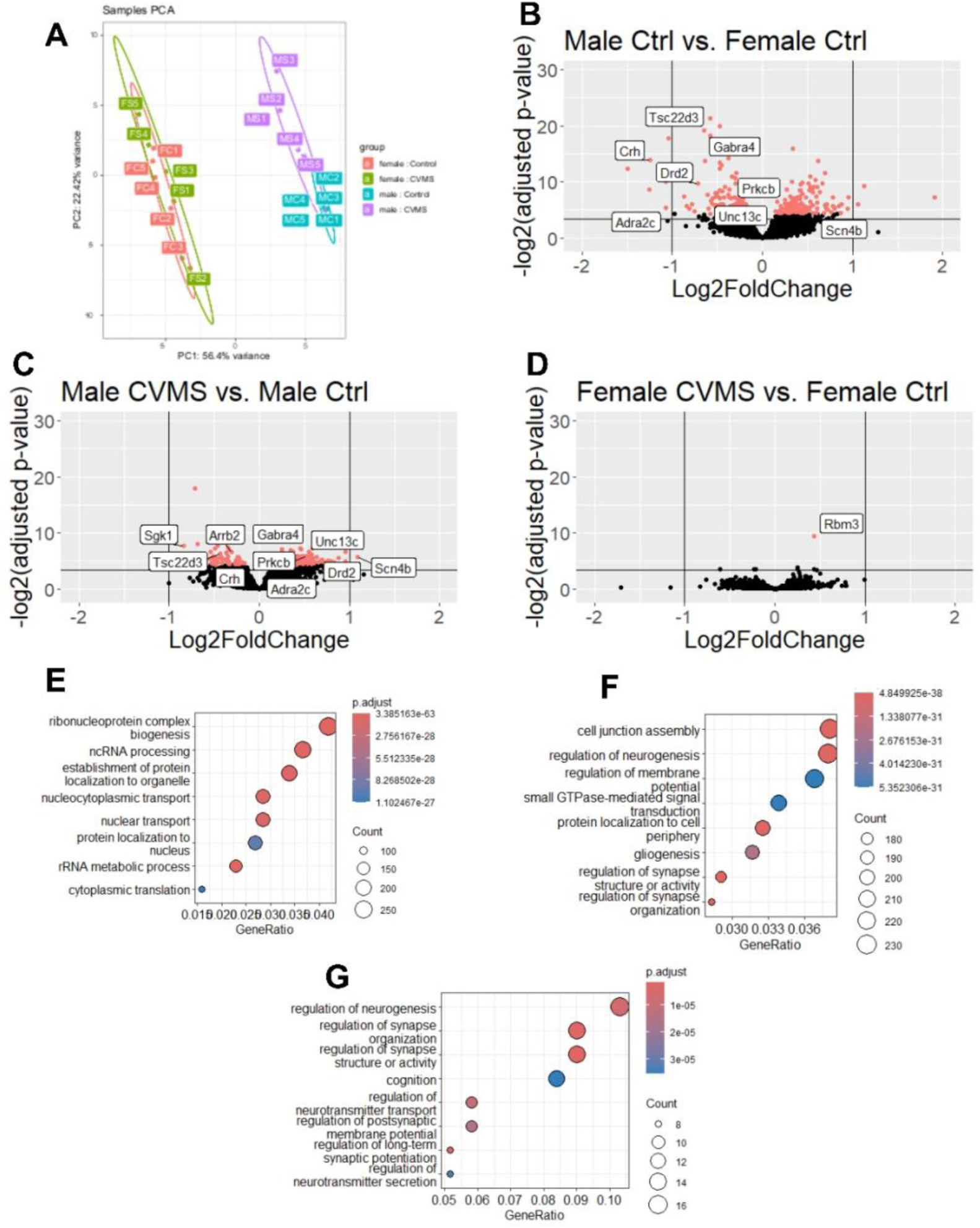
Bulk RNA-Sequencing of the anterodorsal bed nucleus of the stria terminalis. (A) Principal component analysis of all samples. (B) Volcano plot comparing control males and control females. A positive fold change represents a female biased gene while a negative fold change represents a male biased gene. (C & D) Volcano plots comparing control and stressed animals in (C) males and (D) females. A positive fold change represents an upregulation due to stress while a negative fold change represents a downregulation due to stress. (E & F) Gene ontology analyses comparing control males and control females, (E) pathways enriched in control males (F) pathways enriched in control females. (G) Gene ontology analysis comparing control males and stressed males showing pathways upregulated due to stress. A is analyzed by PCAexplorer; B-D are analyzed by DESeq2; and E-F are analyzed by enrichGO. Significant DEGS are highlighted in red in B-D.

Of the genes varying across sex, male biased genes of interest included: *Hcrtr1* (log_2_FC=1.075, adj. p=0.0240) which is the orexin receptor (Scott et al., 2011), *Sgk1* (log_2_FC=1.045, adj. p<0.0001) which modulates neuronal excitability (Lang et al., 2010), *Tac2* (log_2_FC=0.783, adj. p=0.0167) which is the tachykinin/neurokinin B receptor (Al Abed et al., 2021), *Mc4r* (log_2_FC=0.712, adj. p=0.0012) which is a melanocortin receptor (Tao, 2010), *Crh* (log_2_FC=0.572, adj. p=0.0109), *Insyn2a* (log_2_FC=0.280, adj. p=0.0297) which modulates post-synaptic potential (Krueger-Burg et al., 2017), and *Gna13* (log_2_FC=0.198, adj. p=0.0254) which enables dopamine signaling (Yamaguchi et al., 2000). These differences suggest sex-biases in the type and strength of signaling of adBNST neurons as many genes are related to receptors or signaling cascades. A further list of selected male-biased DEGs of interest along with their log_2_FC, adjusted p-value, and functional relevance can be found in Table 1. A list of all male-biased genes can be found in Supplementary Table 2.

**Table 1:**
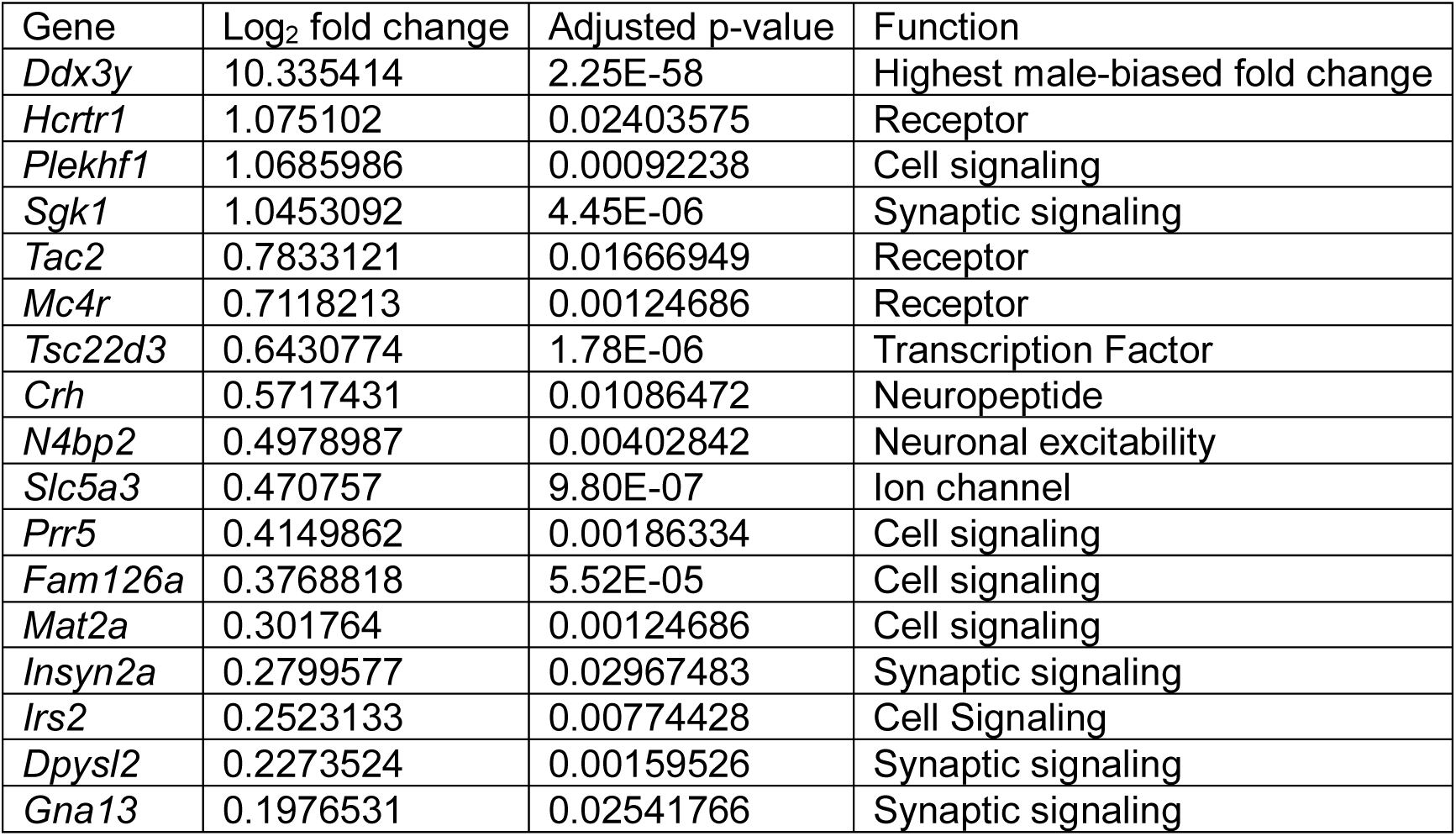
Genes that are male-biased in the adBNST when comparing control males and control females. Genes are listed in order of the log2 fold change. A full list of all male-biased genes can be found in Supplementary Table 1.

Female-biased genes of interest included: *Gsg1l* (log_2_FC=0.603, adj. p=0.0131) and *Cacng3* (log_2_FC=0.428, adj. p=0.0483) which both regulate AMPA receptors (Milstein & Nicoll, 2008; Gu et al., 2016), *Tgfa* (log_2_FC=0.530, adj. p=0.0113) which regulates GnRH signaling and neurogenesis (Jun Ma et al., 1992), *Camk2n1* (log_2_FC=0.443, adj. p=0.0046) which inhibits long-term potentiation (Astudillo et al., 2020), *Cplx1* (log_2_FC=0.281, adj. p=0.0423) and *Cplx2* (log_2_FC=0.350, adj. p=0.0297) which are both important for synaptic vesicle exocytosis (Yoon et al., 2008), *Anks1b* (log_2_FC=0.303, adj. p=0.0405) which modulates synaptic plasticity (Younis et al., 2019), *Comt* (log_2_FC=0.215, adj. p=0.0433) which degrades catecholamines (Harrison & Tunbridge, 2008), and *Slc1a2* (log_2_FC=0.181, adj. p=0.0397) which is a glutamate transporter (Bjørnsen et al., 2014). This suggests a difference in both the signaling and synaptic plasticity in the adBNST between males and females. A further list of selected female-biased DEGs of interest along with their log_2_FC, adjusted p-value, and functional relevance can be found in Table 2. A list of all female-biased genes can be found in Supplementary Table 3.

**Table 2:** Genes that are female-biased in the adBNST when comparing control males and control females. Genes are listed in order of the log2 fold change. A full list of all female-biased genes can be found in Supplementary Table 2.

Female Biased Genes
| Gene | Log<sub>2</sub> fold change | Adjusted p-value | Function |
| --- | --- | --- | --- |
| *Xist* | 12.0753291 | 4.74E-247 | Highest female-biased fold change |
| *Scn4b* | 1.06144811 | 0.01592177 | Ion channel |
| *Unc13c* | 0.69298312 | 0.0003445 | Synaptic signaling |
| *Gsg1l* | 0.60262271 | 0.01313623 | Synaptic signaling |
| *Rgs7bp* | 0.56399579 | 0.00181925 | Synaptic signaling |
| *Tgfa* | 0.52973038 | 0.01129631 | Growth factor |
| *Pcp4* | 0.51704562 | 0.00270595 | Synaptic signaling |
| *Kcnab1* | 0.4574967 | 0.01979391 | Ion channel |
| *Dgkb* | 0.45423177 | 0.02057618 | Synaptic signaling |
| *Camk2n1* | 0.44342939 | 0.00457921 | Synaptic signaling |
| *Cacng3* | 0.42791418 | 0.04830886 | Synaptic signaling |
| *Pcsk2* | 0.41706433 | 0.00572449 | Synaptic signaling |
| *Gabra4* | 0.40890884 | 0.00790706 | Receptor |
| *Gpr158* | 0.39768693 | 0.01796807 | Receptor |
| *Spock3* | 0.38562695 | 0.00673688 | Calcium regulation |
| *Lrrtm3* | 0.36543401 | 0.01125902 | Synaptic signaling |
| *Cplx2* | 0.34982939 | 0.02967483 | Synaptic signaling |
| *Neto2* | 0.32994357 | 0.01532972 | Synaptic signaling |
| *Cdh8* | 0.32613862 | 0.00786683 | Cellular adhesion |
| *Anks1b* | 0.30290019 | 0.04046966 | Synaptic signaling |
| *Adcy1* | 0.29883662 | 0.01387116 | Synaptic signaling |
| *Cacnb4* | 0.28553624 | 0.04816953 | Ion channel |
| *Cplx1* | 0.28103691 | 0.04227382 | Synaptic signaling |
| *Tmem132b* | 0.26624608 | 0.00249415 | Receptor |
| *Comt* | 0.21502622 | 0.04332065 | Synaptic signaling |
| *Gpm6b* | 0.19125504 | 0.03356488 | Synaptic signaling |
| *Slc1a2* | 0.18134176 | 0.03972835 | Transporter |

Comparing control females and stressed females, only 1 gene, *Rbm3*, was found to be differentially regulated and only 6 further genes were trending (Fig. 2d). *Rbm3* was upregulated due to stress (log_2_FC=0.436, adj p<0.0001). Comparing control males to stressed males, 218 genes were found to be significantly differentially expressed with a further 301 trending (Fig. 2c). 161 of the significant genes were found to be upregulated due to stress whereas 57 were downregulated due to stress.

Of the genes altered by stress in males, 217 were differentially regulated with 57 being downregulated by stress and 160 upregulated. Those downregulated by stress included: *Sgk1* (log_2_FC=-0.842, adj. p=0.0050) (Lang et al., 2010), *Slc1a6* (log_2_FC=-0.552, adj. p=0.0232) which is a glutamate transporter (Gegelashvili et al., 2000), *Crh* (log_2_FC=-0.485, adj. p=0.0420), *Vamp1* (log_2_FC=-0.424, adj. p=0.0196) which is important for synaptic vesicle exocytosis (Liu et al., 2011), *Nts* (log_2_FC=-0.409, adj. p=0.0295) which enables neuropeptide receptor binding (Mustain et al., 2011), *Arrb2* (log_2_FC=-0.313, adj. p=0.0348) which modulates β-adrenergic and dopamine signaling (Beaulieu et al., 2005), and *Sorcs3* (log_2_FC=-0.225, adj. p=0.0232) which regulates long-term synaptic depression (Breiderhoff et al., 2013). Those upregulated due to stress included: *Plxnd1* (log_2_FC=0.695, adj. p=0.0263) which is involved in synapse assembly (Wang et al., 2015), *Drd2* (log_2_FC=0.650, adj. p=0.0390) which is a dopamine receptor (Noble, 2000), *Unc13c* (log_2_FC=0.580, adj. p=0.0147) which is involved in glutamatergic transmission (Padmanarayana et al., 2021), *Adra2c* (log_2_FC=0.530, adj. p=0.0484) which is an adrenergic receptor (Bylund, 1992), *Tgfa* (log_2_FC=0.459, adj. p=0.0431) (Jun Ma et al., 1992), *Gabra4* (log_2_FC=0.455, adj. p=0.0067) which is a subunit of the GABA receptor (Mulligan et al., 2012), *Camk2n1* (log_2_FC=0.413, adj. p=0.0196) (Astudillo et al., 2020), *Cplx2* (log_2_FC=0.380, adj, p=0.0230) (Yoon et al., 2008), and *Gpr158* (log_2_FC=0.361, adj. p=0.0431) which is a g-protein coupled receptor that has been implicated in depression (Sutton et al., 2018). Taken together, this suggests that stress causes major changes in synaptic signaling in the adBNST in males. A further list of selected DEGs of interest that are downregulated or upregulated due to stress along with their log_2_FC, adjusted p-value, and functional relevance can be found in Tables 3 and 4 respectively. A list of all stress-regulated genes can be found in Supplementary Table 4.

**Table 3:** Selected genes that are downregulated by stress in the adBNST in males. Genes are listed in order of the log2 fold change. A full list of all genes regulated by stress in males can be found in Supplementary Table 3.

| Gene | Log<sub>2</sub> fold change | Adjusted p-value | Function |
| --- | --- | --- | --- |
| *Sgk1* | -0.8415739 | 0.00501306 | Most downregulated gene |
| *Slc1a6* | -0.5518564 | 0.02320938 | Transporter |
| *Akap12* | -0.5109548 | 0.01745418 | Synaptic signaling |
| *Crh* | -0.4851822 | 0.04204631 | Neuropeptide |
| *Tsc22d3* | -0.4394077 | 0.01335128 | Transcription factor |
| *Vamp1* | -0.4242631 | 0.01961082 | Synaptic signaling |
| *Nts* | -0.4087929 | 0.0294921 | Neuropeptide |
| *Slc5a3* | -0.3474649 | 0.00607686 | Ion channel |
| *Otof* | -0.3466157 | 0.04284925 | Synaptic signaling |
| *Prr5* | -0.3443065 | 0.01854398 | Cell signaling |
| *Alk* | -0.3421647 | 0.03482863 | Synaptic signaling |
| *Arrb2* | -0.3132115 | 0.01075937 | Synaptic signaling |
| *Epha5* | -0.2674514 | 0.04311296 | Receptor |
| *Grk3* | -0.2649624 | 0.04374024 | Synaptic signaling |
| *Sema4a* | -0.2401795 | 0.04735542 | Synaptic signaling |
| *Sorcs3* | -0.22523 | 0.02320938 | Synaptic signaling |
| *Dpysl2* | -0.1670872 | 0.04284925 | Synaptic signaling |

**Table 4:** Selected genes that are upregulated by stress in the adBNST in males. Genes are listed in order of the log2 fold change. A full list of all genes regulated by stress in males can be found in Supplementary Table 3.

| Gene | Log<sub>2</sub> fold change | Adjusted p-value | Function |
| --- | --- | --- | --- |
| *Scn4b* | 1.08954915 | 0.01961082 | Most upregulated gene |
| *Rasd2* | 0.87684047 | 0.02308458 | Synaptic signaling |
| *Ppp1r1b* | 0.7098293 | 0.03707918 | Cell signaling |
| *Plxnd1* | 0.69541227 | 0.02628802 | Synaptic signaling |
| *Itpr1* | 0.6703348 | 0.03462583 | Receptor |
| *Drd2* | 0.65004132 | 0.03902923 | Receptor |
| *Ano2* | 0.61227969 | 0.00607686 | Ion channel |
| *Gng7* | 0.59981204 | 0.03850164 | Receptor |
| *Musk* | 0.5834368 | 0.02217479 | Synaptic signaling |
| *Unc13c* | 0.58046104 | 0.01465707 | Synaptic signaling |
| *Slc18a3* | 0.57407102 | 0.0426947 | Transporter |
| *Adra2c* | 0.52976636 | 0.04836964 | Receptor |
| *Penk* | 0.50832335 | 0.03386401 | Neuropeptide |
| *Ddn* | 0.50724373 | 0.01961082 | Synaptic signaling |
| *Rgs7bp* | 0.50103361 | 0.01817878 | Synaptic signaling |
| *Dapp1* | 0.48901824 | 0.01854398 | Cell signaling |
| *Kcnab1* | 0.48442864 | 0.01961082 | Ion channel |
| *Lzts3* | 0.4828807 | 0.03043355 | Synaptic signaling |
| *Kcna4* | 0.47734195 | 0.04524505 | Ion channel |
| *Tgfa* | 0.45877183 | 0.04311296 | Growth factor |
| *Gabra4* | 0.45498929 | 0.006678 | Receptor |
| *Filip1* | 0.45307766 | 0.03310177 | Synaptic signaling |
| *Mctp1* | 0.43760904 | 0.04204631 | Synaptic signaling |
| *Epha7* | 0.42772808 | 0.04204631 | Receptor |
| *Camk2n1* | 0.41345675 | 0.0195909 | Synaptic signaling |
| *Pcsk2* | 0.40659244 | 0.01681495 | Protein modification |
| *Shank3* | 0.39997231 | 0.04311296 | Synaptic signaling |
| *Cplx2* | 0.38001612 | 0.02296941 | Synaptic signaling |
| *Gpr158* | 0.36139748 | 0.04311296 | Receptor |
| *Spock3* | 0.34781457 | 0.02857924 | Calcium regulation |
| *Igsf11* | 0.29617933 | 0.03138945 | Synaptic signaling |
| *Adcy1* | 0.26192307 | 0.04311296 | Synaptic signaling |
| *Tmem123b* | 0.25428606 | 0.01075937 | Receptor |
| *Frmpd4* | 0.23296982 | 0.01681495 | Synaptic signaling |

Comparing control males and control females for gene ontology enrichment, pathways that were male biased in controls were mostly related to RNA processing and protein localization (Fig. 2e) whereas pathways upregulated in control females are related to neuronal activity, such as neurogenesis and regulation of the synapse (Fig, 2f). Comparing control males to stressed males, pathways that are upregulated due to stress are mostly related to synaptic activity and neurotransmitters (Fig. 2g). There were no pathways that were significantly downregulated due to stress in males.

## Electrophysiology

### M-current

In examination of stress-induced changes in excitability in the adBNST from CRH-expressing neurons, we found no differences in the resting membrane potential between the experimental groups. Control males had an average RMP of −69.2 ± 2.2 mV, stressed males had an average of −69.0 ± 1.9 mV, control females had an average of −68.7 ± 1.9 mV, and stressed females had an average of −67.7 ± 2.3 mV (Fig, 3b). We also found no differences in the change in Rin resistance due to XE-991, the KCNQ channel blocker. The average change in Rin in control males was −0.39 ± 0.18 pA, in stressed males was −0.38 ± 0.09 pA, in control females was −0.43 ± 0.11 pA, and in stressed females was −0.22 ± 0.14 pA (Fig. 3c).

**Figure 3:**
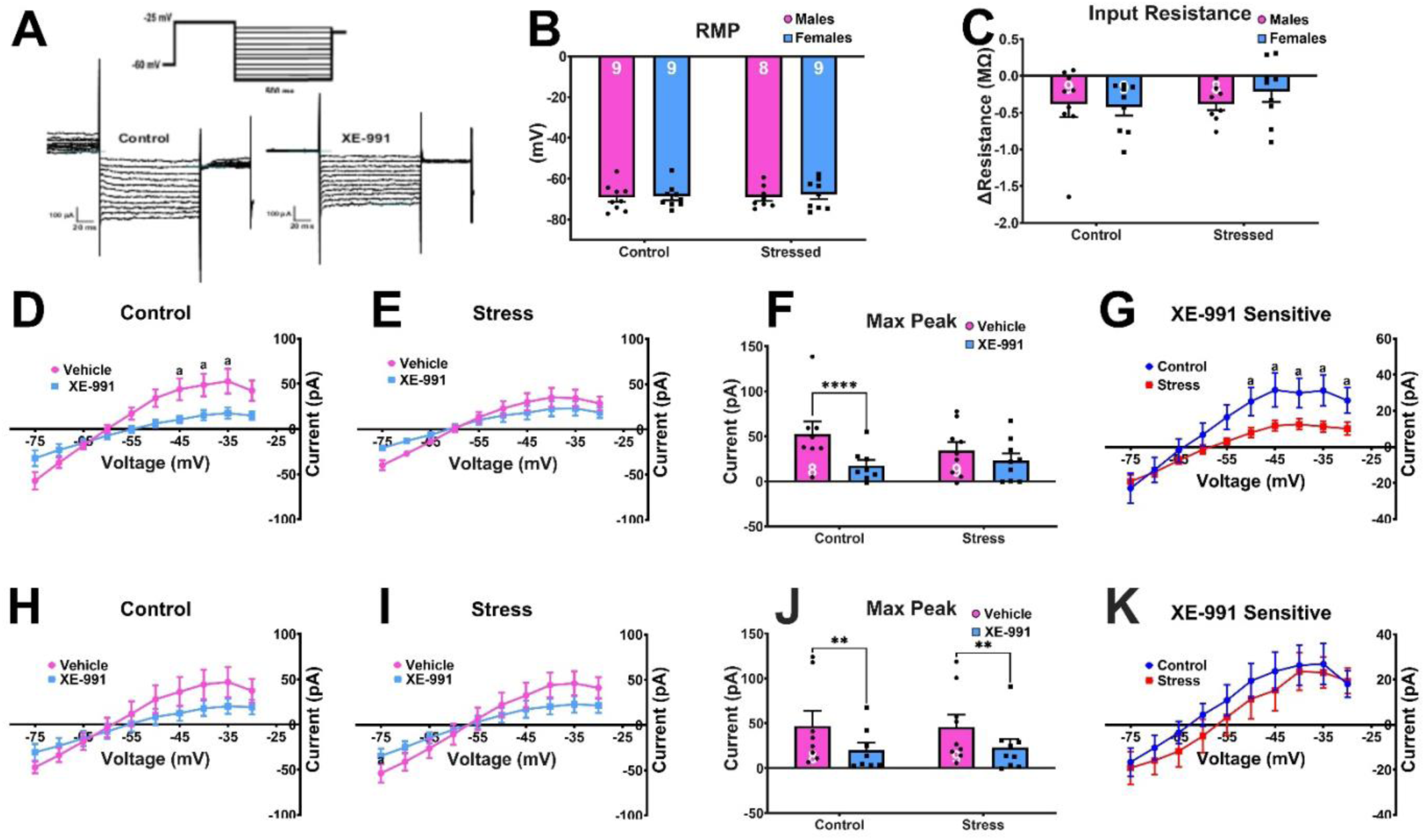
Recordings of M-current activity of BNST Corticotropin Releasing Factor-expressing neurons from stressed and non-stressed mice. (A) M-current protocol and representative traces. (B) Resting membrane potential of the neurons. (C) Change in input resistance after perfusion with XE-991. I-V plots from −75 to −30 mV after XE-991 perfusion in (D) Control male mice and (E) CVMS-exposed male mice. (F) Current at max peak in male mice. (G) I-V plot of XE-991 sensitive current in male mice. (H-K) Control and CVMS-exposed female mice. Data are presented as mean +/-SEM. (a=0.5-0.1) (**=0.01-0.001, ****<0.0001).

We next examined the M-current and found an effect of CVMS in the males, as we previously found (Hu 2020), but not CRH neurons from female mice. XE-991 perfusion suppressed the M-current activity in control male mice but not stressed male mice (Fig. 3d & e). The max peak, defined as the strength of the current in −35mV trace, in CRH neurons from control males decreased after exposure to XE-991 (p<0.0001) but not in CRH neurons from stressed males (Fig. 3f). Additionally, the traces of the XE-991 sensitive current were decreased in CRH neurons from stressed males in comparison to control males (Fig. 3g).

CRH neurons from female mice did not show changes in the M-current after exposure to XE-991 in either control or stressed groups (Fig. 3h & i). However, max peak was significantly reduced in both control (p=0.0052) and stressed (p=0.0092) groups after XE-991 perfusion, suggesting that the M-current is active in these neurons. Further, no difference in the XE-991 sensitive current in CRH neurons from female mice were found due to CVMS.

### EPSCs

We next examined the influence of CVMS on EPSC in both sexes. Previous studies in our lab suggest that CVMS augments EPSCs in male mice (Hu 2020). In the current study, CVMS in male mice did not have change the number of EPSCs, the amplitude of the EPSCs, or the area under the curve (AUC) of the EPSCs. Control males in vehicle conditions had an average of 159.4 ± 47.5 EPSCs events while stressed males had an average of 145 ± 43.8 EPSCs. For amplitude, CRH neurons from control males in vehicle, EPSCs had an average amplitude of - 8.65 ± 1.0 pA whereas neurons from stressed males had an average of −9.8 ± 1.0 pA. For AUC, CRH neurons from control males in vehicle, EPSCs had an average AUC of −82.7 ± 12.6 pA*ms whereas neurons from stressed males had an average of −71.2 ± 15.3 pA*ms.

After initial recording of EPSCs, we perfused each CRH neuron with Stressin 1, an agonist of CRHR1 receptors, to determine if CRH may be acting in a feedback loop in these neuroins, either in an autocrine or paracrine fashion.. After perfusion with Stressin I, CRH neurons from control males had an average of 131.5 ± 42.5 EPSCs and neurons from stressed males had an average of 115 ± 21.7 (Fig, 4a). CRH neurons from control male EPSCs had an average amplitude of −9.0 ± 0.8 pA whereas neurons from stressed males had an average of - 10.3 ± 0.9 pA. EPSCs from control males had an average AUC of −68.3 ± 2.9 pA*ms and neurons from stressed males had an average of –78.0 ± 12.6 pA*ms (Fig. 4c).

**Figure 4:**
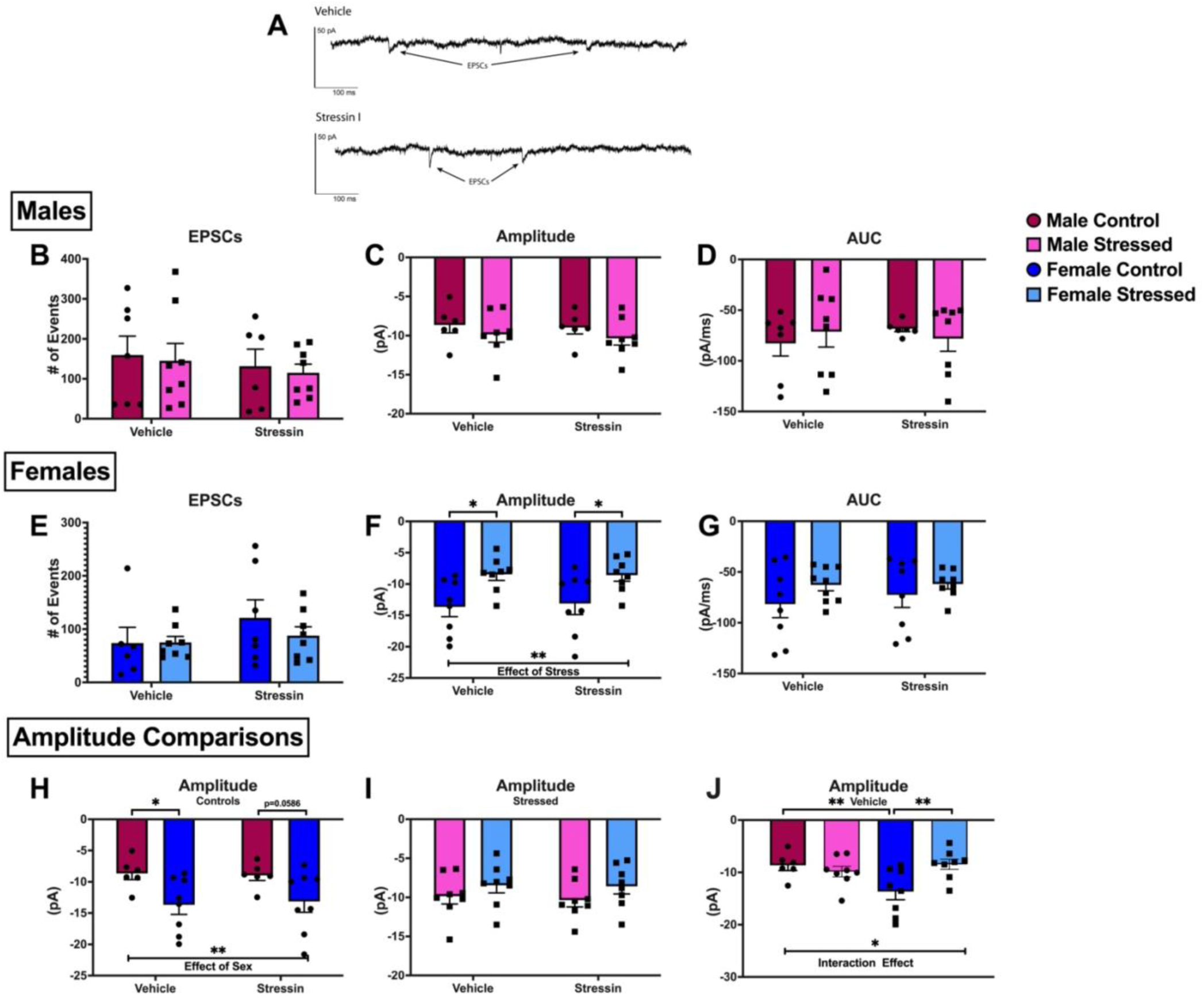
Recordings of EPSCs in CRF-expressing neurons of the adBNST from control and stressed mice. (A) Representative traces of current recordings for EPSCs in vehicle and after exposure to Stressin I. Number of EPSCs detected in a 5-minute voltage-clamp recording in vehicle and after exposure to Stressin I in (B) male and (E) female mice. Average amplitude of EPSCs detected in (C) male and (F) female mice. Average area under the curve of EPSCs detected in (D) male and (G) female mice. (H) Comparison of average amplitude in males and females in control mice. (I) Comparison of average amplitude in male and female mice in stressed mice. (J) Comparison of average amplitude between males and females and control and stressed when cells are in vehicle conditions. Data is presented as mean +/-SEM. Analyzed via two-way ANOVA (*=0.05-0.01, **=0.01-0.001).

CRH neurons from female mice showed differences in the amplitude of their EPSCs due to stress. CRH neurons from control females in vehicle conditions had an average of 73.8 ± 29.5 EPSCs while neurons from stressed females had an average of 75.0 ± 11.2 EPSCs. CRH neurons from control females in vehicle, EPSCs had an average amplitude of −13.7 ± 1.6 pA whereas neurons from stressed females had an average of −8.5 ± 1.0 pA, this was significant with p=0.0116. For AUC, CRH neurons from control females in vehicle, EPSCs had an average AUC of −81.7 ± 13.3 pA*ms whereas neurons from stressed females had an average of −62.7± 5.7 pA*ms.

After perfusion with Stressin I, CRH neurons from control females had an average of 121.1 ± 33.7 EPSCs while CRH neurons from stressed females had an average of 87.8 EPSCs ± 16.6 (Fig. 4d). The neurons from control female EPSCs had an average amplitude of −13.1 ± 1.8 pA whereas neurons from stressed females had an average of −8.6 ± 1.0 pA (p=0.0259, Fig. 4e), with a main effect of stress in the amplitude of EPSCs (F(1,28)=12.77, p=0.0013). CRH neurons from control females had an average AUC of −72.4 ± 12.6 pA*ms and neurons from stressed females had an average AUC of –61.9 ± 4.9 pA*ms (Fig. 4f).

When comparing males and females, we do see differences in the amplitude of the EPSCs. Both in vehicle and Stressin I, CRH neurons from females had significantly lower amplitudes to their EPSCs with a main effect of sex of F(1,24)=9.587, p=0.0049, Fig. 4g). We do not observe differences in the stressed animals, suggesting that CVMS is causing the female CRH-expressing neurons to behave similarly to the males (Fig. 4h). This is further supported by the interaction effect in Fig. 4i which suggest that the EPSCs in CRH neurons from females and males are responding differently to stress (F(1,26)=7.114, p=0.0130).

## DISCUSSION

Our previous research using CVMS suggested that the transcriptome of the adBNST was not affected by stress nor did it affect the M-current in NPY neurons of the adBNST (Degroat et al., 2024). This seemingly contradicted our hypothesis that the BNST was a major processing center for chronic stress. However, these current finding necessitate further research into the BNST and its relations to stress signaling, mood disorders, and sex variability. In contrast to our past research, this study suggests that there are transient changes in the transcriptome due to stress in males but not females, and also suggests that the neurons in the BNST and their signaling properties are altered by stress, but in differing ways across the sexes.

We found an abundance of genes that were differentially expressed in the males and females and also regulated by stress in the males. Interestingly, we found no such differences in the females, already suggesting that stress is affecting this area differently in males and females. The lack of response to CVMS in females also suggests that the adBNST is an important mediator of stress in males but not in females and that the major central stress regulating region in females may be elsewhere. This hypothesis is further supported by the fact that presynaptic EPSCs were affected by stress in the females but the not males.

Alternatively, this data could suggest that CVMS is not effective on females in general. As we did not see differences due to stress in final cumulative BW gain or adBNST transcriptome, this suggests there may be a resistance to CVMS. Historically, literature has been contradictory in how sex relates to sensitivity of CMVS in mice (Hodes, 2018). This issue is further compounded by the limited research of CVMS in mice compared to rats, the limited research of CVMS in females, and potential strain differences which also show differing sensitivities to stress (Mineur et al., 2006). However, our previous publication showed that females were sensitive to our CVMS paradigm, but that sensitivity was dependent on the estrous cycles (Degroat et al., 2024). Further research should control for estrous stage in the RNA-sequencing, along with electrophysiology tests in differing estrous stages in order to test this. We believe this, along with our data showing increased EPSCs strength in stressed females, suggests the need for further research in the effectiveness of CVMS in females.

In terms of sex differences in the transcriptome of the adBNST, the data shows that male-biased genes are related to transcriptional regulation while female-biased genes are related to synaptic processing, suggesting that the BNST in the females may be more plastic, especially at their synapse. For example, female-biased genes include *Gsg1l* and *Cacng3* which both regulate AMPA signaling. *Gsg1l* acts to negatively regulate AMPA receptors by slowing the deactivation and desensitization of AMPA receptors (Greger et al., 2017). *Cacng3* is a transmembrane AMPA regulatory protein (TARP) which is a family of proteins that positively regulate AMPA receptors. TARPs act by promoting AMPA receptor trafficking to the synapse and stabilization in the postsynaptic density (Coombs & Cull-Candy, 2009). With the transcripts of both negatively and positively regulating proteins being female-biased, suggests that the neurons in the female adBNST are more primed for synaptic plasticity than males. This hypothesis is further supported by other genes involved in synaptic plasticity and axonogenesis, such as *Anks2b* and *Pcp4*, are female-biased.

However, while the female adBNST may be primed for synaptic plasticity, the data suggests that only the transcriptome of the male adBNST is sensitive to stress. Of the genes that are downregulated by stress, many are related to synaptic signaling. For example, many genes are signaling molecules, their receptors, or regulators of their receptors, such as *Crh*, corticotropin releasing hormone, *Slc1a6*, a glutamate transporter (Gegelashvili et al., 2000), and *Arrb2*, and inhibitor of arrestin receptors(Beaulieu et al., 2005). Additionally, some are related to synaptic maintenance and signaling in general, such as *Vamp1*, which is involved in synaptic vesicle exocytosis (Liu et al., 2011), *Otof*, which is involved in synaptic vesicle priming (Johnson & Chapman, 2010), and *Dpysl2*, which is involved in axon growth (Inagaki et al., 2001).

Upregulated genes, also fall a similar pattern of affecting synaptic transmission and catecholamine signaling. Examples include: *Drd2,* a dopamine receptor (Noble, 2000), *Adra2c*, an adrenergic receptor (Bylund, 1992), and *Camk2n1* and *Igsf11*, which are both involved in long-term potentiation (Jang et al., 2016; Astudillo et al., 2020). This suggests that chronic stress causes major changes in the regulation of the synapse in the adBNST of males.

However, one must ask if this is truly a reflection of chronic stress or acute stress. Our previous publication showed no transcriptomic differences due to CVMS 18 h post-stress (Degroat et al., 2024) but in the current study, we found over 200 DEGs in the males 1 h post-stress. However, the transcriptional effects of chronic stress did lead to sustained (17-24 h post-stress) neurophysiological outcomes in males. It is possible that our results are due specifically to chronic stress and the transcriptome is so transient that difference are lost quickly when there is no stress, or it could be equally possible that this is due to the mild acute stress. One study using chronic mild stress paradigms found sustained differences in the nucleus accumbensm24 h after the last stressor (Ma et al., 2019). Two other studies found stress sensitive genes in the amygdala, hippocampus, and hypothalamus after chronic stress but waited to collect tissue until after multiple days of behavior testing (Karisetty et al., 2017; Yang et al., 2023). Comparisons between chronic stress studies can be difficult though as the chronic stress paradigms often differ significantly between labs and the transcriptome of different brain regions or cell types may have differing reaction times to stress. Additonally, Karisetty et al. and Yang et al. used behavior tests to ensure the mice expressed stress-induced behavioral changes before transcriptomic analysis, which we did not do, further complicating comparisons. Further experiments may be necessary, such as measuring the adBNST transcriptome in samples collected one hour after the typical last stressor (1 h wet bedding) without the chronic stress paradigm.

As the transcriptome data suggest that only the male adBNST is sensitive to stress, this is also supported by chronic stress affecting the M-current in males but not females. Chronic stress significantly decreased the strength of the M-current in males, supporting our previously published findings (Hu et al., 2020). The M-current is an intrinsic property of the neuron and isproduced by voltage-gated K^+^ channel, the KCNQ channels. This current is inhibitory and is significant as it is one of the few currents that is active at physiologically relevant voltages. This weakening of the M-current by stress infers that these neurons are more likely to fire, leading to increased release of CRH or GABA and further supports our hypothesis that CVMS directly affects the adBNST neurons in males but not females.

The females suggest there may be different areas of the brain that are more important for chronic stress processing. As the transcriptome data suggests, the adBNST in females is not sensitive to CVMS as the only gene affected by it was *Rbm3*, a gene that is related to cellular stress (Hu et al., 2022). Additionally, the electrophysiology data supports examining other regions critical to stress or steroid hormone signaling as we saw significant differences in the females to EPSCs. EPSCs being an extrinsic property shows that neurons other than the CRF-expressing adBNST neurons are being affected by stress and we are seeing that downstream through their synaptic transmission (Egli & Winder, 2003). Furthermore, our data is supported by the lack of response to CRH receptor activation either in the M-current or EPSCs, demonstrating that adBNST CRH neurons do not respond to other CRH neurons. However, this does not preclude other types of neurons in the BNST causing these changes, One potential site is the posterior BNST which expresses high levels of steroid receptors and send projections to the adBNST (Becker et al., 2008; Sun et al., 2023).

Our data suggests that chronic stress is processed differently in the brains of male and female mice. However, surprisingly, it also suggests that they’re not only processed differently, but potentially processed by separate areas of the brain. Further research is needed to elucidate which other areas are important and how these differences in signaling is related to the development of mood disorders. Additionally, this experiment only used one type of stress paradigm. It is well known that different types of stress are processed differently, thus further studies are also needed to be done to determine how the processing of psychosocial stress may differ from physiological stress for example.

## REFERENCES

Al Abed, A. S., Reynolds, N. J., & Dehorter, N. (2021). A Second Wave for the Neurokinin Tac2 Pathway in Brain Research. Biological Psychiatry, 90(3), 156–164. 10.1016/j.biopsych.2021.02.016

Albert, K. M., & Newhouse, P. A. (2019). Estrogen, Stress, and Depression: Cognitive and Biological Interactions. Annu Rev Clin Psychol, 15, 399–423. 10.1146/annurev-clinpsy-050718-095557

Astudillo, D., Karmelic, D., Casas, B. S., Otmakhov, N., Palma, V., & Sanhueza, M. (2020). CaMKII inhibitor 1 (CaMK2N1) mRNA is upregulated following LTP induction in hippocampal slices. Synapse, 74(10), e22158. 10.1002/syn.22158

Beaulieu, J.-M., Sotnikova, T. D., Marion, S., Lefkowitz, R. J., Gainetdinov, R. R., & Caron, M. G. (2005). An Akt/β-Arrestin 2/PP2A Signaling Complex Mediates Dopaminergic Neurotransmission and Behavior. Cell, 122(2), 261–273. 10.1016/j.cell.2005.05.012

Becker, J. A. J., Befort, K., Blad, C., Filliol, D., Ghate, A., Dembele, D., Thibault, C., Koch, M., Muller, J., Lardenois, A., Poch, O., & Kieffer, B. L. (2008). Transcriptome analysis identifies genes with enriched expression in the mouse central extended amygdala. Neuroscience, 156(4), 950–965. 10.1016/j.neuroscience.2008.07.070

Bjørnsen, L. P., Hadera, M. G., Zhou, Y., Danbolt, N. C., & Sonnewald, U. (2014). The GLT-1 (EAAT2; slc1a2) glutamate transporter is essential for glutamate homeostasis in the neocortex of the mouse. Journal of Neurochemistry, 128(5), 641–649. 10.1111/jnc.12509

Breiderhoff, T., Christiansen, G. B., Pallesen, L. T., Vaegter, C., Nykjaer, A., Holm, M. M., Glerup, S., & Willnow, T. E. (2013). Sortilin-related receptor SORCS3 is a postsynaptic modulator of synaptic depression and fear extinction. PLoS One, 8(9), e75006. 10.1371/journal.pone.0075006

Bromet, E., Andrade, L. H., Hwang, I., Sampson, N. A., Alonso, J., de Girolamo, G., de Graaf, R., Demyttenaere, K., Hu, C., Iwata, N., Karam, A. N., Kaur, J., Kostyuchenko, S., Lépine, J. P., Levinson, D., Matschinger, H., Mora, M. E., Browne, M. O., Posada-Villa, J., … Kessler, R. C. (2011). Cross-national epidemiology of DSM-IV major depressive episode. BMC Med, 9, 90. 10.1186/1741-7015-9-90

Bylund, D. B. (1992). Subtypes of alpha 1- and alpha 2-adrenergic receptors. Faseb j, 6(3), 832–839. 10.1096/fasebj.6.3.1346768

Coombs, I. D., & Cull-Candy, S. G. (2009). Transmembrane AMPA receptor regulatory proteins and AMPA receptor function in the cerebellum. Neuroscience, 162(3), 656–665. 10.1016/j.neuroscience.2009.01.004

Degroat, T. J., Wiersielis, K., Denney, K., Kodali, S., Daisey, S., Tollkuhn, J., Samuels, B. A., & Roepke, T. A. (2024). Chronic stress and its effects on behavior, RNA expression of the bed nucleus of the stria terminalis, and the M-current of NPY neurons. Psychoneuroendocrinology, 161, 106920. 10.1016/j.psyneuen.2023.106920

Egli, R. E., & Winder, D. G. (2003). Dorsal and Ventral Distribution of Excitable and Synaptic Properties of Neurons of the Bed Nucleus of the Stria Terminalis. Journal of Neurophysiology, 90(1), 405–414. 10.1152/jn.00228.2003

Freeman, E. W. (2003). Premenstrual syndrome and premenstrual dysphoric disorder: definitions and diagnosis11Adapted from the symposium on Premenstrual Syndrome and Premenstrual Dysphoric Disorders, July 17, 2000, Rhodes, Greece. Psychoneuroendocrinology, 28, 25–37. 10.1016/S0306-4530(03)00099-4

Gegelashvili, G., Dehnes, Y., Danbolt, N. C., & Schousboe, A. (2000). The high-affinity glutamate transporters GLT1, GLAST, and EAAT4 are regulated via different signalling mechanisms. Neurochemistry International, 37(2), 163–170. 10.1016/S0197-0186(00)00019-X

Greger, I. H., Watson, J. F., & Cull-Candy, S. G. (2017). Structural and Functional Architecture of AMPA-Type Glutamate Receptors and Their Auxiliary Proteins. Neuron, 94(4), 713–730. 10.1016/j.neuron.2017.04.009

Gu, X., Mao, X., Lussier, M. P., Hutchison, M. A., Zhou, L., Hamra, F. K., Roche, K. W., & Lu, W. (2016). GSG1L suppresses AMPA receptor-mediated synaptic transmission and uniquely modulates AMPA receptor kinetics in hippocampal neurons. Nature Communications, 7(1), 10873. 10.1038/ncomms10873

Hammack, S. E., Todd, T. P., Kocho-Schellenberg, M., & Bouton, M. E. (2015). Role of the bed nucleus of the stria terminalis in the acquisition of contextual fear at long or short context-shock intervals. Behav Neurosci, 129(5), 673–678. 10.1037/bne0000088

Harrison, P. J., & Tunbridge, E. M. (2008). Catechol-O-Methyltransferase (COMT): A Gene Contributing to Sex Differences in Brain Function, and to Sexual Dimorphism in the Predisposition to Psychiatric Disorders. Neuropsychopharmacology, 33(13), 3037–3045. 10.1038/sj.npp.1301543

Herman, J. P., Ostrander, M. M., Mueller, N. K., & Figueiredo, H. (2005). Limbic system mechanisms of stress regulation: hypothalamo-pituitary-adrenocortical axis. Prog Neuropsychopharmacol Biol Psychiatry, 29(8), 1201–1213. 10.1016/j.pnpbp.2005.08.006

Hodes, G. E. (2018). A primer on sex differences in the behavioral response to stress. Current Opinion in Behavioral Sciences, 23, 75–83. 10.1016/j.cobeha.2018.03.012

Hu, P., Liu, J., Yasrebi, A., Gotthardt, J. D., Bello, N. T., Pang, Z. P., & Roepke, T. A. (2016). Gq Protein-Coupled Membrane-Initiated Estrogen Signaling Rapidly Excites Corticotropin-Releasing Hormone Neurons in the Hypothalamic Paraventricular Nucleus in Female Mice. Endocrinology, 157(9), 3604–3620. 10.1210/en.2016-1191

Hu, P., Maita, I., Phan, M. L., Gu, E., Kwok, C., Dieterich, A., Gergues, M. M., Yohn, C. N., Wang, Y., Zhou, J. N., Qi, X. R., Swaab, D. F., Pang, Z. P., Lucassen, P. J., Roepke, T. A., & Samuels, B. A. (2020). Early-life stress alters affective behaviors in adult mice through persistent activation of CRH-BDNF signaling in the oval bed nucleus of the stria terminalis. Transl Psychiatry, 10(1), 396. 10.1038/s41398-020-01070-3

Hu, Y., Liu, Y., Quan, X., Fan, W., Xu, B., & Li, S. (2022). RBM3 is an outstanding cold shock protein with multiple physiological functions beyond hypothermia. J Cell Physiol, 237(10), 3788–3802. 10.1002/jcp.30852

Inagaki, N., Chihara, K., Arimura, N., Ménager, C., Kawano, Y., Matsuo, N., Nishimura, T., Amano, M., & Kaibuchi, K. (2001). CRMP-2 induces axons in cultured hippocampal neurons. Nature Neuroscience, 4(8), 781–782. 10.1038/90476

Jang, S., Oh, D., Lee, Y., Hosy, E., Shin, H., van Riesen, C., Whitcomb, D., Warburton, J. M., Jo, J., Kim, D., Kim, S. G., Um, S. M., Kwon, S.-k., Kim, M.-H., Roh, J. D., Woo, J., Jun, H., Lee, D., Mah, W., … Kim, E. (2016). Synaptic adhesion molecule IgSF11 regulates synaptic transmission and plasticity. Nature Neuroscience, 19(1), 84–93. 10.1038/nn.4176

Johnson, C. P., & Chapman, E. R. (2010). Otoferlin is a calcium sensor that directly regulates SNARE-mediated membrane fusion. Journal of Cell Biology, 191(1), 187–197. 10.1083/jcb.201002089

Jun Ma, Y., Junier, M.-P., Costa, M. E., & Ojeda, S. R. (1992). Transforming growth factor-α gene expression in the hypothalamus is developmentally regulated and linked to sexual maturation. Neuron, 9(4), 657–670. 10.1016/0896-6273(92)90029-D

Karisetty, B. C., Khandelwal, N., Kumar, A., & Chakravarty, S. (2017). Sex difference in mouse hypothalamic transcriptome profile in stress-induced depression model. Biochemical and Biophysical Research Communications, 486(4), 1122–1128. 10.1016/j.bbrc.2017.04.005

Krueger-Burg, D., Papadopoulos, T., & Brose, N. (2017). Organizers of inhibitory synapses come of age. Current Opinion in Neurobiology, 45, 66–77. 10.1016/j.conb.2017.04.003

Lai, C.-H. (2011). Major Depressive Disorder: Gender Differences in Symptoms, Life Quality, and Sexual Function. Journal of Clinical Psychopharmacology, 31(1). https://journals.lww.com/psychopharmacology/fulltext/2011/02000/major_depress ive_disorder gender_differences_in.8.aspx

Lang, F., Strutz-Seebohm, N., Seebohm, G., & Lang, U. E. (2010). Significance of SGK1 in the regulation of neuronal function. J Physiol, 588(Pt 18), 3349–3354. 10.1113/jphysiol.2010.190926

Li, J., Liu, F., Liu, Z., Li, M., Wang, Y., Shang, Y., & Li, Y. (2024). Prevalence and associated factors of depression in postmenopausal women: a systematic review and meta-analysis. BMC Psychiatry, 24(1), 431. 10.1186/s12888-024-05875-0

Liu, Y., Sugiura, Y., & Lin, W. (2011). The role of Synaptobrevin1/VAMP1 in Ca2+-triggered neurotransmitter release at the mouse neuromuscular junction. The Journal of Physiology, 589(7), 1603–1618. 10.1113/jphysiol.2010.201939

Ma, K., Zhang, H., Wei, G., Dong, Z., Zhao, H., Han, X., Song, X., Zhang, H., Zong, X., Baloch, Z., & Wang, S. (2019). Identification of key genes, pathways and miRNA/mRNA regulatory networks of CUMS-induced depression in nucleus accumbens by integrated bioinformatics analysis. Neuropsychiatric Disease and Treatment, 15, 685–700. 10.2147/NDT.S200264

Milstein, A. D., & Nicoll, R. A. (2008). Regulation of AMPA receptor gating and pharmacology by TARP auxiliary subunits. Trends Pharmacol Sci, 29(7), 333–339. 10.1016/j.tips.2008.04.004

Mineur, Y. S., Belzung, C., & Crusio, W. E. (2006). Effects of unpredictable chronic mild stress on anxiety and depression-like behavior in mice. Behavioural Brain Research, 175(1), 43–50. 10.1016/j.bbr.2006.07.029

Mulligan, M. K., Wang, X., Adler, A. L., Mozhui, K., Lu, L., & Williams, R. W. (2012). Complex control of GABA(A) receptor subunit mRNA expression: variation, covariation, and genetic regulation. PLoS One, 7(4), e34586. 10.1371/journal.pone.0034586

Mustain, W. C., Rychahou, P. G., & Evers, B. M. (2011). The role of neurotensin in physiologic and pathologic processes. Curr Opin Endocrinol Diabetes Obes, 18(1), 75–82. 10.1097/MED.0b013e3283419052

Noble, E. P. (2000). The DRD2 gene in psychiatric and neurological disorders and its phenotypes. Pharmacogenomics, 1(3), 309–333. 10.1517/14622416.1.3.309

O’Hara, M. W., & McCabe, J. E. (2013). Postpartum depression: current status and future directions. Annu Rev Clin Psychol, 9, 379–407. 10.1146/annurev-clinpsy-050212-185612

Padmanarayana, M., Liu, H., Michelassi, F., Li, L., Betensky, D., Dominguez, M. J., Sutton, R. B., Hu, Z., & Dittman, J. S. (2021). A unique C2 domain at the C terminus of Munc13 promotes synaptic vesicle priming. Proc Natl Acad Sci U S A, 118(11). 10.1073/pnas.2016276118

Poe, G. R., Foote, S., Eschenko, O., Johansen, J. P., Bouret, S., Aston-Jones, G., Harley, C. W., Manahan-Vaughan, D., Weinshenker, D., Valentino, R., Berridge, C., Chandler, D. J., Waterhouse, B., & Sara, S. J. (2020). Locus coeruleus: a new look at the blue spot. Nature Reviews Neuroscience, 21(11), 644–659. 10.1038/s41583-020-0360-9

Scott, M. M., Marcus, J. N., Pettersen, A., Birnbaum, S. G., Mochizuki, T., Scammell, T. E., Nestler, E. J., Elmquist, J. K., & Lutter, M. (2011). Hcrtr1 and 2 signaling differentially regulates depression-like behaviors. Behavioural Brain Research, 222(2), 289–294. 10.1016/j.bbr.2011.02.044

Sun, Y., Zweifel, L. S., Holmes, T. C., & Xu, X. (2023). Whole-brain input mapping of the lateral versus medial anterodorsal bed nucleus of the stria terminalis in the mouse. Neurobiology of Stress, 23, 100527. 10.1016/j.ynstr.2023.100527

Sutton, L. P., Orlandi, C., Song, C., Oh, W. C., Muntean, B. S., Xie, K., Filippini, A., Xie, X., Satterfield, R., Yaeger, J. D. W., Renner, K. J., Young, S. M., Jr., Xu, B., Kwon, H., & Martemyanov, K. A. (2018). Orphan receptor GPR158 controls stress-induced depression. Elife, 7. 10.7554/eLife.33273

Tao, Y. X. (2010). The melanocortin-4 receptor: physiology, pharmacology, and pathophysiology. Endocr Rev, 31(4), 506–543. 10.1210/er.2009-0037

Wang, F., Eagleson, K. L., & Levitt, P. (2015). Positive regulation of neocortical synapse formation by the Plexin-D1 receptor. Brain Research, 1616, 157–165. 10.1016/j.brainres.2015.05.005

Yamaguchi, Y., Katoh, H., Yasui, H., Aoki, J., Nakamura, K., & Negishi, M. (2000). Gα12 and Gα13 Inhibit Ca2+ -Dependent Exocytosis Through Rho/Rho-Associated Kinase-Dependent Pathway. Journal of Neurochemistry, 75(2), 708–717. 10.1046/j.1471-4159.2000.0750708.x

Yang, S., Yi, L., Xia, X., Chen, X., Hou, X., Zhang, L., Yang, F., Liao, J., Han, Z., & Fu, Y. (2023). Transcriptome comparative analysis of amygdala-hippocampus in depression: A rat model induced by chronic unpredictable mild stress (CUMS). Journal of Affective Disorders, 334, 258–270. 10.1016/j.jad.2023.04.074

Yoon, T.-Y., Lu, X., Diao, J., Lee, S.-M., Ha, T., & Shin, Y.-K. (2008). Complexin and Ca2+ stimulate SNARE-mediated membrane fusion. Nature Structural & Molecular Biology, 15(7), 707–713. 10.1038/nsmb.1446

Younis, R. M., Taylor, R. M., Beardsley, P. M., & McClay, J. L. (2019). The ANKS1B gene and its associated phenotypes: focus on CNS drug response. Pharmacogenomics, 20(9), 669–669–684. 10.2217/pgs-2019-0015

