## Supplementary Table 1. for "Chronic variable mild stress alters the transcriptome and signaling properties of the anterodorsal bed nuceleus of the stria terminalis in a sex-dependent manner"

**Appendix 1**

**Supplementary Table 1.**

All outliers for the EPSCs analyses.

| | Male Control | Male Stressed | Female Control | Female Stressed |
|---|---|---|---|---|
| Sample Size | n=7 | n=8 | n-8 | n=9 |
| Pre Stressin 1 | | | | |
| EPSCs | | | 1 outlier | 1 outlier |
| Amplitude | 1 outlier | | | 1 outlier |
| AUC | | | | |
| Post Stressin 1 | | | | |
| EPSCs | 1 outlier | | | 1 outlier |
| Amplitude | 1 outlier | | | 1 outlier |
| AUC | 1 outlier | | | 1 outlier |

**Supplementary Table 2.**

All male-biased genes in the adBNST when comparing control males and control females.

Genes are ordered by largest Log2FoldChange.

| Male-Biased Genes | | | | |
|---|---|---|---|---|
| | baseMean | log2FoldChange | pvalue | padj |
| Kdm5d | 313.865675 | 10.7627754 | 3.57E-33 | 1.16E-29 |
| Ddx3y | 713.891281 | 10.3354142 | 3.47E-62 | 2.25E-58 |
| Eif2s3y | 363.521715 | 10.0848461 | 9.24E-41 | 4.00E-37 |
| Uty | 178.72564 | 9.36500624 | 1.10E-29 | 2.39E-26 |
| Hspa1b | 61.6883226 | 1.49496076 | 3.26E-07 | 0.00019765 |
| Sytl4 | 238.726532 | 1.25207753 | 8.64E-06 | 0.00249415 |
| Hspa1a | 57.7918478 | 1.24383076 | 8.27E-08 | 6.71E-05 |
| Hcrtr1 | 41.0816708 | 1.075102 | 0.00026704 | 0.02403575 |
| Plekhf1 | 57.5137158 | 1.06859863 | 2.06E-06 | 0.00092238 |
| Sgk1 | 1207.86131 | 1.04530924 | 4.46E-09 | 4.45E-06 |
| Hif3a | 163.701719 | 0.86771931 | 3.27E-05 | 0.00634566 |
| Fosl2 | 487.230053 | 0.83752257 | 0.00020828 | 0.01979391 |
| Map3k15 | 62.9062024 | 0.81101179 | 0.00032145 | 0.02633192 |
| Tac2 | 643.068423 | 0.78331208 | 0.00014537 | 0.01666949 |
| Sap30 | 45.6794006 | 0.74964624 | 0.00057705 | 0.0386324 |
| Mc4r | 71.5726854 | 0.71182126 | 3.30E-06 | 0.00124686 |
| Tsc22d3 | 1004.29073 | 0.64307735 | 1.50E-09 | 1.78E-06 |

| | | | | |
|---|---|---|---|---|
| Arrdc2 | 155.914339 | 0.63980035 | 9.95E-08 | 7.27E-05 |
| Zfp57 | 236.193702 | 0.61087105 | 2.10E-05 | 0.00462005 |
| Klf2 | 123.79884 | 0.60187156 | 0.00032236 | 0.02633192 |
| Dnajb1 | 860.991174 | 0.58158816 | 2.93E-09 | 3.17E-06 |
| Gpr161 | 319.80106 | 0.57987358 | 2.65E-10 | 3.82E-07 |
| Vipr2 | 87.038359 | 0.57745533 | 0.00094124 | 0.04929357 |
| Crh | 280.43959 | 0.57174307 | 7.45E-05 | 0.01086472 |
| Nfkbia | 214.035946 | 0.56108185 | 0.00011168 | 0.01400883 |
| Arid5a | 123.112074 | 0.55301004 | 4.53E-05 | 0.00754187 |
| Gpr165 | 701.19167 | 0.53353318 | 0.00018205 | 0.01861515 |
| P4ha2 | 199.371047 | 0.51757837 | 1.64E-05 | 0.00388272 |
| Trib1 | 141.374419 | 0.5148999 | 3.09E-05 | 0.00616568 |
| Ldlr | 314.943272 | 0.50197757 | 0.00017891 | 0.01844204 |
| N4bp2 | 616.830125 | 0.4978987 | 1.74E-05 | 0.00402842 |
| Zmym1 | 892.238884 | 0.48767263 | 0.00062144 | 0.03972835 |
| Sh2b2 | 113.511294 | 0.47872821 | 4.07E-05 | 0.00697506 |
| Spry4 | 254.201511 | 0.47780269 | 3.35E-07 | 0.00019765 |
| Ddit4 | 423.555928 | 0.47555606 | 1.05E-05 | 0.00274112 |
| Ccdc117 | 368.832337 | 0.47395907 | 1.38E-07 | 9.40E-05 |
| Slc5a3 | 1408.77153 | 0.47075702 | 7.55E-10 | 9.80E-07 |
| Tiparp | 422.658461 | 0.4690772 | 5.21E-05 | 0.00786683 |
| Dgkk | 892.173333 | 0.46663766 | 0.00015016 | 0.01666949 |
| Bcl6 | 508.398831 | 0.45076848 | 1.95E-07 | 0.00012673 |
| Lsm11 | 208.058195 | 0.44568455 | 1.27E-05 | 0.00317243 |
| Col25a1 | 1927.53375 | 0.43175066 | 0.00014998 | 0.01666949 |
| Zfp658 | 144.695089 | 0.42809673 | 3.45E-05 | 0.00659426 |
| Prr5 | 249.217694 | 0.41498621 | 6.03E-06 | 0.00186334 |
| Rbm12b2 | 269.400813 | 0.40112985 | 0.00043864 | 0.03165026 |
| Pde11a | 126.520836 | 0.39257477 | 0.00072876 | 0.04321953 |
| Adcy6 | 936.251092 | 0.39212771 | 0.00035222 | 0.02772483 |
| Txnip | 526.56357 | 0.38998856 | 0.00027548 | 0.02403575 |
| Cpne7 | 1531.05236 | 0.38313704 | 0.00038919 | 0.02967483 |
| Zfp948 | 252.111766 | 0.380244 | 2.32E-05 | 0.00502284 |
| Fam126a | 685.728361 | 0.37688176 | 6.38E-08 | 5.52E-05 |
| Ahsa2 | 470.588729 | 0.37203213 | 3.71E-05 | 0.00673688 |
| Rbm3 | 530.177425 | 0.36973606 | 9.10E-06 | 0.00256914 |
| Clk1 | 2179.30816 | 0.35959326 | 0.0004159 | 0.03051822 |
| Sik1 | 388.442828 | 0.35839563 | 0.00020701 | 0.01979391 |
| Arid5b | 613.423799 | 0.34489548 | 4.65E-05 | 0.00765048 |
| Tbkbp1 | 852.688219 | 0.33801029 | 9.63E-07 | 0.00048563 |
| Bag3 | 168.413651 | 0.33748594 | 0.0008158 | 0.04606778 |

| | | | | |
|---|---|---|---|---|
| Edrf1 | 446.834172 | 0.32830424 | 0.00016409 | 0.01761285 |
| Jmjd6 | 338.340008 | 0.32527486 | 0.00088017 | 0.04816953 |
| Gprasp2 | 5134.69828 | 0.31293778 | 0.00048314 | 0.03356488 |
| Lonrf3 | 519.380441 | 0.31180232 | 3.97E-05 | 0.00697506 |
| Yeats2 | 738.435148 | 0.31003569 | 0.00084458 | 0.04667816 |
| Mycl | 279.364975 | 0.30672204 | 0.00046453 | 0.03315021 |
| Mat2a | 4018.10958 | 0.30176403 | 3.29E-06 | 0.00124686 |
| Camk1g | 961.156379 | 0.30164437 | 0.0006761 | 0.04131053 |
| Adipor2 | 1249.32552 | 0.30053782 | 1.30E-06 | 0.0006267 |
| Pprc1 | 563.725549 | 0.29200666 | 0.00010957 | 0.01395222 |
| Nufip2 | 1386.95208 | 0.29198126 | 4.96E-05 | 0.00775531 |
| Hmgcr | 1356.89133 | 0.28774765 | 6.17E-07 | 0.0003445 |
| Insyn2a | 318.304679 | 0.27995768 | 0.00038958 | 0.02967483 |
| Zfp516 | 621.376584 | 0.27624674 | 0.00067205 | 0.04131053 |
| Klf15 | 560.455306 | 0.26852085 | 0.00017891 | 0.01844204 |
| Tnpo1 | 1052.56989 | 0.25940283 | 8.19E-05 | 0.01125902 |
| Trp53inp1 | 371.328761 | 0.25685816 | 0.00090082 | 0.04816953 |
| F3 | 692.43367 | 0.2566067 | 0.00093582 | 0.04920813 |
| Hap1 | 12300.6765 | 0.25656174 | 0.00089222 | 0.04816953 |
| Irs2 | 1858.78089 | 0.25231327 | 4.87E-05 | 0.00774428 |
| Tmtc3 | 383.810887 | 0.24558638 | 0.00079805 | 0.04556562 |
| Ocrl | 2104.52895 | 0.24033603 | 1.81E-05 | 0.00412654 |
| D10Wsu102e | 472.197574 | 0.23993081 | 0.00028441 | 0.02446286 |
| Tub | 4152.24894 | 0.2344995 | 0.00016159 | 0.01748913 |
| Klhl42 | 518.913249 | 0.23423765 | 0.00027759 | 0.02403575 |
| Hsph1 | 5691.10901 | 0.23259837 | 2.96E-05 | 0.00600142 |
| Tmed8 | 1512.42555 | 0.23231241 | 7.87E-05 | 0.01123311 |
| Ddx50 | 975.806886 | 0.2313309 | 0.00060827 | 0.03930418 |
| Tro | 6953.55976 | 0.23112676 | 3.77E-06 | 0.0013244 |
| Ccdc6 | 1562.27942 | 0.23052026 | 0.00013568 | 0.01600163 |
| Brd3 | 1344.78478 | 0.22967069 | 4.12E-05 | 0.00697506 |
| Dpysl2 | 2963.42914 | 0.22735241 | 4.67E-06 | 0.00159526 |
| Cdc37l1 | 2710.75331 | 0.2269346 | 0.00050403 | 0.03482106 |
| Uhrf2 | 1061.74831 | 0.22568696 | 0.00038403 | 0.02967483 |
| Tra2b | 947.682113 | 0.22273511 | 7.82E-05 | 0.01123311 |
| Dnajb4 | 1345.17996 | 0.22168347 | 0.00011791 | 0.01458433 |
| Cpeb2 | 1691.78369 | 0.2208364 | 0.00059896 | 0.03910705 |
| Mfhas1 | 675.98687 | 0.21906661 | 0.00090216 | 0.04816953 |
| Trmt1l | 538.32267 | 0.21873494 | 0.00071107 | 0.04275621 |
| Slc2a1 | 1546.03723 | 0.2186266 | 0.00059633 | 0.03910705 |
| Zfp260 | 776.080862 | 0.21464647 | 0.00017345 | 0.01831557 |

| | | | | |
|---|---|---|---|---|
| Ubxn7 | 1132.21631 | 0.21360246 | 0.0008034 | 0.04556562 |
| Crkl | 1122.28365 | 0.21099846 | 4.85E-05 | 0.00774428 |
| Gda | 8326.32809 | 0.20455492 | 0.00082699 | 0.04609873 |
| Scyl2 | 812.028116 | 0.20209674 | 0.00048302 | 0.03356488 |
| Asb1 | 849.623339 | 0.2011093 | 0.00080095 | 0.04556562 |
| Ets2 | 1932.81741 | 0.20052722 | 0.00078644 | 0.04540152 |
| Pnmal2 | 12524.3381 | 0.19930623 | 0.00080273 | 0.04556562 |
| Gna13 | 1303.83601 | 0.19765307 | 0.0003058 | 0.02541766 |
| Gprasp1 | 17586.4325 | 0.19560643 | 0.00037625 | 0.02926219 |
| Spast | 1128.5175 | 0.19434082 | 0.0002999 | 0.02541766 |
| Narf | 753.130209 | 0.19015844 | 0.00093204 | 0.04920813 |
| Zbtb11 | 1099.41175 | 0.18665979 | 0.00013875 | 0.01609041 |
| Fam53c | 1028.87707 | 0.17842409 | 0.00021563 | 0.02020035 |
| Zfp354c | 977.976933 | 0.1687568 | 0.00040405 | 0.03010228 |
| Ppm1f | 1383.32012 | 0.15712786 | 0.00027236 | 0.02403575 |
| Zfp664 | 2207.91562 | 0.14076615 | 0.00027646 | 0.02403575 |
| Fam91a1 | 1874.53781 | 0.13524484 | 0.00077375 | 0.04506487 |

**Supplementary Table 3.**

All female-biased genes in the adBNST when comparing control males and control females.

Genes are ordered by largest Log2FoldChange.

| Female-Biased Genes | | | | |
|---|---|---|---|---|
| | baseMean | log2FoldChange | pvalue | padj |
| Xist | 9946.78343 | 12.0753291 | 3.65E-251 | 4.74E-247 |
| Clspn | 62.617626 | 1.9041297 | 4.01E-05 | 0.00697506 |
| Tmem200b | 59.9974352 | 1.1261665 | 5.24E-06 | 0.0017443 |
| Scn4b | 6254.74511 | 1.06144811 | 0.0001327 | 0.01592177 |
| Gask1b | 79.6305789 | 0.95481065 | 3.59E-05 | 0.00673688 |
| Cd4 | 439.480907 | 0.94228861 | 0.00033674 | 0.02683189 |
| 4833415N18Rik | 68.9506882 | 0.87535905 | 0.00014708 | 0.01666949 |
| Mafa | 50.2308284 | 0.86405683 | 0.00089269 | 0.04816953 |
| Rasd2 | 3721.54175 | 0.84646321 | 0.00020879 | 0.01979391 |
| Nexn | 518.833199 | 0.80402224 | 0.00027387 | 0.02403575 |
| Acvr1c | 443.132635 | 0.80253063 | 0.00034103 | 0.02700764 |
| Rbp4 | 90.9499562 | 0.78444261 | 0.00021619 | 0.02020035 |
| Ky | 62.6435898 | 0.77605482 | 3.66E-05 | 0.00673688 |
| Pde10a | 9334.17076 | 0.76092682 | 0.00071583 | 0.042844 |
| Eif2s3x | 489.513507 | 0.72684231 | 1.40E-25 | 2.60E-22 |
| Adora2a | 1822.62492 | 0.71986689 | 0.00042396 | 0.03093457 |

| | | | | |
|---|---|---|---|---|
| Hpca | 6606.28765 | 0.70804058 | 0.00014973 | 0.01666949 |
| Asic4 | 558.293383 | 0.70033713 | 0.00030601 | 0.02541766 |
| Unc13c | 4779.04151 | 0.69298312 | 6.37E-07 | 0.0003445 |
| BC031361 | 94.2074769 | 0.68942842 | 0.00018346 | 0.01861515 |
| Gpr88 | 6841.39947 | 0.68846436 | 0.000773 | 0.04506487 |
| Coch | 366.592142 | 0.66273191 | 2.83E-06 | 0.00115043 |
| Gpd1 | 640.914132 | 0.63748073 | 1.49E-05 | 0.0036429 |
| Exph5 | 138.840846 | 0.61713085 | 1.01E-07 | 7.27E-05 |
| Rgs4 | 4436.5296 | 0.60738166 | 5.68E-06 | 0.00181925 |
| Ano3 | 4180.3679 | 0.60558332 | 0.00036303 | 0.02840369 |
| Gsg1l | 1174.95258 | 0.60262271 | 9.91E-05 | 0.01313623 |
| Pcp4l1 | 3367.86876 | 0.59834828 | 2.94E-05 | 0.00600142 |
| Foxp1 | 2824.99842 | 0.5981692 | 0.00089484 | 0.04816953 |
| Cd59a | 121.523053 | 0.58812116 | 1.62E-06 | 0.0007504 |
| Rgs7bp | 3794.26515 | 0.56399579 | 5.74E-06 | 0.00181925 |
| Lpar6 | 59.9931693 | 0.56158231 | 0.00078652 | 0.04540152 |
| Arhgdib | 276.121723 | 0.56060834 | 0.00015853 | 0.0173026 |
| Rasgrp1 | 5737.86512 | 0.55586988 | 1.52E-05 | 0.0036656 |
| Rasgef1b | 745.3761 | 0.54707562 | 9.88E-06 | 0.00270595 |
| Kdm6a | 825.094839 | 0.53721943 | 7.04E-19 | 1.14E-15 |
| AI593442 | 2622.75571 | 0.53184068 | 9.72E-07 | 0.00048563 |
| Tgfa | 1343.72403 | 0.52973038 | 8.35E-05 | 0.01129631 |
| Fbxo32 | 504.815281 | 0.52722545 | 6.19E-06 | 0.00186934 |
| Ddx3x | 5534.85391 | 0.52659085 | 2.37E-32 | 6.16E-29 |
| Krt10 | 139.290911 | 0.52649306 | 0.00022843 | 0.02104179 |
| Dlgap3 | 3911.55079 | 0.51902344 | 0.00041422 | 0.03051822 |
| Pcp4 | 3482.84278 | 0.51704562 | 1.00E-05 | 0.00270595 |
| Ldlrad4 | 413.729684 | 0.50654749 | 3.36E-06 | 0.00124686 |
| Robo2 | 1608.92985 | 0.49041426 | 0.00028736 | 0.02455413 |
| Thsd7a | 1786.15232 | 0.48599821 | 0.00023314 | 0.02132415 |
| Pdzd2 | 3031.6821 | 0.48251353 | 0.00032629 | 0.02648617 |
| Cep128 | 140.390873 | 0.48242538 | 0.00052684 | 0.03601805 |
| Pde7b | 3222.90215 | 0.47859721 | 0.00062272 | 0.03972835 |
| Dhrsx | 146.559599 | 0.4784513 | 4.14E-05 | 0.00697506 |
| Jcad | 2867.16044 | 0.47802479 | 0.00048193 | 0.03356488 |
| Them6 | 314.121702 | 0.46457448 | 0.00052781 | 0.03601805 |
| Kcnab1 | 3579.66627 | 0.4574967 | 0.00020493 | 0.01979391 |
| Dgkb | 6501.68119 | 0.45423177 | 0.00022179 | 0.02057618 |
| Pcdhga9 | 243.887569 | 0.45401861 | 0.00033337 | 0.02683189 |
| Sema3e | 529.296069 | 0.44726371 | 0.00017876 | 0.01844204 |
| Lmo2 | 427.837108 | 0.44710247 | 0.0006718 | 0.04131053 |

| | | | | |
|---|---|---|---|---|
| Caln1 | 1326.56257 | 0.44382393 | 0.00065123 | 0.04046966 |
| Camk2n1 | 10281.142 | 0.44342939 | 2.04E-05 | 0.00457921 |
| Cldn5 | 330.814345 | 0.4398186 | 2.68E-06 | 0.00112475 |
| Prkcb | 17960.1132 | 0.43871952 | 0.00082307 | 0.04609873 |
| Gpr26 | 533.919605 | 0.43049148 | 0.00087322 | 0.04805658 |
| P2ry13 | 212.697281 | 0.42970242 | 0.00043162 | 0.03131772 |
| Pcdhb14 | 168.47996 | 0.42912015 | 0.00012048 | 0.01476169 |
| Cacng3 | 1177.49858 | 0.42791418 | 0.00091128 | 0.04830886 |
| Klf3 | 297.899956 | 0.42380074 | 1.06E-05 | 0.00274112 |
| St6galnac5 | 933.579235 | 0.42274656 | 3.62E-06 | 0.00130699 |
| Pcsk2 | 2957.93364 | 0.41706433 | 2.69E-05 | 0.00572449 |
| Hmgcs2 | 580.525274 | 0.41094517 | 1.13E-05 | 0.0028813 |
| Gabra4 | 1964.11491 | 0.40890884 | 5.30E-05 | 0.00790706 |
| Gpr158 | 2826.27972 | 0.39768693 | 0.00016878 | 0.01796807 |
| Ppp3ca | 22331.1738 | 0.39496613 | 0.00026871 | 0.02403575 |
| Atp2b1 | 11481.7976 | 0.39193699 | 0.00060695 | 0.03930418 |
| Map6d1 | 797.398915 | 0.38700556 | 0.00065018 | 0.04046966 |
| Spock3 | 3993.33259 | 0.38562695 | 3.73E-05 | 0.00673688 |
| Apcdd1 | 357.27027 | 0.37774167 | 0.00011217 | 0.01400883 |
| Zc3h6 | 453.358738 | 0.37556183 | 5.14E-05 | 0.00786683 |
| Nrde2 | 316.022626 | 0.36977515 | 4.89E-05 | 0.00774428 |
| Lrrtm3 | 1362.41036 | 0.36543401 | 8.20E-05 | 0.01125902 |
| Man1a | 1613.45779 | 0.36114507 | 0.00063191 | 0.04003512 |
| Cobl | 1533.43713 | 0.35860332 | 0.00030725 | 0.02541766 |
| Lrrtm1 | 893.17542 | 0.35247978 | 0.00010171 | 0.01334346 |
| Cplx2 | 18164.0609 | 0.34982939 | 0.0003907 | 0.02967483 |
| Coq8a | 459.561165 | 0.3483215 | 0.0004056 | 0.03010228 |
| Rapgef4 | 8141.75275 | 0.34169433 | 0.00013362 | 0.01592177 |
| Kdm5c | 1565.56479 | 0.33854307 | 1.74E-08 | 1.62E-05 |
| Zfp365 | 5459.89321 | 0.33520129 | 0.00059919 | 0.03910705 |
| Jup | 992.188509 | 0.33325842 | 0.00018782 | 0.01887419 |
| St8sia3 | 5411.20199 | 0.33249753 | 0.00019467 | 0.019301 |
| Lrrc8b | 1787.8312 | 0.33184691 | 8.05E-05 | 0.01125902 |
| Calb1 | 4796.27589 | 0.33165539 | 0.00013676 | 0.01600163 |
| Cxcl12 | 784.551971 | 0.33115793 | 2.21E-06 | 0.00095724 |
| Neto2 | 1880.71499 | 0.32994357 | 0.00012629 | 0.01532972 |
| Sp2 | 401.318852 | 0.32646262 | 0.00064108 | 0.04041922 |
| Cdh8 | 1074.41649 | 0.32613862 | 5.21E-05 | 0.00786683 |
| Scara3 | 460.806469 | 0.32478271 | 8.55E-05 | 0.0114463 |
| Smarcd1 | 2664.26402 | 0.31715448 | 0.00056121 | 0.03796366 |
| Cntn4 | 523.981115 | 0.31179778 | 2.75E-05 | 0.00575998 |

| | | | | |
|---|---|---|---|---|
| Otud7a | 428.509101 | 0.31166679 | 0.00018892 | 0.01887419 |
| Ctsz | 448.736577 | 0.31054533 | 0.00052968 | 0.03601805 |
| Clvs1 | 475.944804 | 0.30792148 | 3.22E-05 | 0.00633913 |
| Tgfbr2 | 313.005769 | 0.30369438 | 0.00030157 | 0.02541766 |
| Anks1b | 6115.24599 | 0.30290019 | 0.00064826 | 0.04046966 |
| Adcy1 | 2300.47846 | 0.29883662 | 0.00010787 | 0.01387116 |
| Mapk4 | 1311.42041 | 0.29351909 | 0.00090494 | 0.04816953 |
| Zdhhc14 | 823.340459 | 0.29264333 | 0.0001534 | 0.01688405 |
| Tomm70a | 5526.20344 | 0.28637685 | 0.00072036 | 0.04291779 |
| Cacnb4 | 3900.61992 | 0.28553624 | 0.00088373 | 0.04816953 |
| Cap1 | 4622.11206 | 0.28154263 | 0.00058738 | 0.03910705 |
| Cplx1 | 3936.54471 | 0.28103691 | 0.00069979 | 0.04227382 |
| Ptprb | 917.078421 | 0.27515952 | 0.00047782 | 0.03356488 |
| Plppr5 | 1021.88421 | 0.27069354 | 0.00033503 | 0.02683189 |
| Tmem132b | 2428.61821 | 0.26624608 | 8.59E-06 | 0.00249415 |
| Adora1 | 1290.53865 | 0.26534171 | 0.00026776 | 0.02403575 |
| Cnnm2 | 668.184637 | 0.26403253 | 0.00059883 | 0.03910705 |
| Lrrc8d | 852.972827 | 0.2634994 | 0.00019855 | 0.01953611 |
| Gpr155 | 1548.8982 | 0.2582656 | 0.00068375 | 0.04149795 |
| Prkca | 2843.99487 | 0.25404349 | 6.61E-05 | 0.0097495 |
| St3gal5 | 1258.30182 | 0.25240216 | 0.0001059 | 0.01375473 |
| Zfp36l2 | 581.629725 | 0.25183714 | 0.00040143 | 0.03010228 |
| Maneal | 999.698797 | 0.23146747 | 0.0004595 | 0.03297255 |
| Ap1ar | 1642.75702 | 0.22318751 | 0.00039669 | 0.02995481 |
| Comt | 1220.86868 | 0.21502622 | 0.0007338 | 0.04332065 |
| Galnt13 | 1448.01691 | 0.21177386 | 0.00082668 | 0.04609873 |
| Sorbs1 | 4356.45021 | 0.19948342 | 0.00020237 | 0.01976182 |
| Kif5c | 13562.8885 | 0.19941716 | 0.00075171 | 0.04417738 |
| Acss1 | 694.876901 | 0.19677305 | 0.00067748 | 0.04131053 |
| Gpm6b | 17237.6156 | 0.19125504 | 0.00048326 | 0.03356488 |
| Calm2 | 17891.3581 | 0.18139095 | 0.00083256 | 0.04621065 |
| Slc1a2 | 45912.1177 | 0.18134176 | 0.00062401 | 0.03972835 |
| Slc39a10 | 2439.4721 | 0.16393318 | 8.24E-05 | 0.01125902 |
| Plpp3 | 5609.10734 | 0.13988408 | 0.00056621 | 0.03810305 |

**Supplementary Table 4.**

All genes that were shown to be differentially expressed according to stress in males. A positive Log2FoldChange represents a gene upregulated by stress and a negative Log2FoldChange is downregulated by stress. Genes are ordered by largest Log2FoldChange.

| Male Stress-Sensitve Genes |
|---|

| | baseMean | log2FoldChange | pvalue | padj |
|---|---|---|---|---|
| Scn4b | 6254.74511 | 1.08954915 | 8.71E-05 | 0.01961082 |
| 4833415N18Rik | 68.9506882 | 0.9577981 | 1.68E-05 | 0.01075937 |
| Cd4 | 439.480907 | 0.94956583 | 0.00027929 | 0.03297611 |
| Rasd2 | 3721.54175 | 0.87684047 | 0.00012056 | 0.02308458 |
| Grb7 | 44.854639 | 0.82659543 | 0.0003939 | 0.03680585 |
| Ido1 | 76.8297167 | 0.82346471 | 0.00076203 | 0.04735542 |
| Acvr1c | 443.132635 | 0.79632008 | 0.00034251 | 0.03482863 |
| Gm10754 | 80.0903263 | 0.79265208 | 0.00059143 | 0.04311296 |
| Neu2 | 55.9839698 | 0.76207193 | 0.00020678 | 0.0294921 |
| Pde10a | 9334.17076 | 0.76016846 | 0.00072117 | 0.04688472 |
| Adora2a | 1822.62492 | 0.71969591 | 0.00041188 | 0.0376445 |
| Rbp4 | 90.9499562 | 0.71894869 | 0.00048637 | 0.04054544 |
| Ppp1r1b | 8219.96731 | 0.7098293 | 0.00039968 | 0.03707918 |
| Plxnd1 | 1173.08178 | 0.69541227 | 0.00016076 | 0.02628802 |
| Itpr1 | 9318.96889 | 0.6703348 | 0.00033058 | 0.03462583 |
| Cep128 | 140.390873 | 0.65187627 | 8.54E-07 | 0.00408363 |
| Drd2 | 1528.10511 | 0.65004132 | 0.0004476 | 0.03902923 |
| Lpl | 391.109438 | 0.63821405 | 9.68E-05 | 0.02061329 |
| Ano2 | 352.928772 | 0.61227969 | 2.89E-06 | 0.00607686 |
| Epop | 81.6981674 | 0.60840308 | 0.00036231 | 0.03511653 |
| Ddit4l | 250.643149 | 0.60320895 | 0.00056745 | 0.04284925 |
| Gng7 | 9871.08159 | 0.59981204 | 0.00043577 | 0.03850164 |
| Lmo2 | 427.837108 | 0.59303106 | 3.90E-06 | 0.00633156 |
| Them6 | 314.121702 | 0.59181599 | 5.51E-06 | 0.006678 |
| Musk | 90.106347 | 0.5834368 | 0.00010756 | 0.02217479 |
| Unc13c | 4779.04151 | 0.58046104 | 2.93E-05 | 0.01465707 |
| Ano3 | 4180.3679 | 0.57958241 | 0.00063178 | 0.04435402 |
| Pcp4l1 | 3367.86876 | 0.57846411 | 5.13E-05 | 0.0179986 |
| Arhgap10 | 313.167908 | 0.5758233 | 0.00019569 | 0.02859076 |
| Slc18a3 | 77.4688841 | 0.57407102 | 0.00054513 | 0.0426947 |
| Prkch | 674.168429 | 0.56535933 | 0.00032405 | 0.03449827 |
| Jcad | 2867.16044 | 0.55646121 | 4.53E-05 | 0.01682116 |
| Slc35d3 | 349.784256 | 0.55603804 | 0.00024135 | 0.03043355 |
| Gpr26 | 533.919605 | 0.54558911 | 1.70E-05 | 0.01075937 |
| Crybg3 | 550.625219 | 0.53353244 | 0.00030991 | 0.03382479 |
| Arpp19 | 4228.37299 | 0.53331851 | 3.88E-05 | 0.01681495 |
| Adra2c | 407.772738 | 0.52976636 | 0.0007858 | 0.04836964 |
| Tesc | 457.497495 | 0.52860052 | 0.00041014 | 0.0376445 |
| Fbxl16 | 17854.0175 | 0.52446863 | 9.13E-05 | 0.01975438 |
| Gpd1 | 640.914132 | 0.52205218 | 0.00033853 | 0.03482863 |

| | | | | |
|---|---|---|---|---|
| Prkcb | 17960.1132 | 0.51518027 | 8.48E-05 | 0.01961082 |
| Gcnt2 | 635.055275 | 0.50989039 | 0.00032797 | 0.03462583 |
| Penk | 11088.2925 | 0.50832335 | 0.00031288 | 0.03386401 |
| Rasgrp1 | 5737.86512 | 0.50760405 | 7.59E-05 | 0.01961082 |
| Caln1 | 1326.56257 | 0.50745662 | 8.45E-05 | 0.01961082 |
| Ddn | 10591.0261 | 0.50724373 | 7.64E-05 | 0.01961082 |
| Ier5l | 165.488515 | 0.50379443 | 0.00052558 | 0.04213715 |
| Robo2 | 1608.92985 | 0.50171918 | 0.00018924 | 0.02857924 |
| Rgs7bp | 3794.26515 | 0.50103361 | 5.32E-05 | 0.01817878 |
| Pdzd2 | 3031.6821 | 0.49369713 | 0.00022445 | 0.0294921 |
| Dapp1 | 126.234398 | 0.48901824 | 6.28E-05 | 0.01854398 |
| Kcnab1 | 3579.66627 | 0.48442864 | 7.97E-05 | 0.01961082 |
| Lzts3 | 4609.07739 | 0.4828807 | 0.00023967 | 0.03043355 |
| Pde7b | 3222.90215 | 0.47926952 | 0.00058936 | 0.04311296 |
| Kcna4 | 805.726219 | 0.47734195 | 0.00064973 | 0.04524505 |
| Ankrd63 | 2318.36407 | 0.47441759 | 0.00020649 | 0.0294921 |
| 2010300C02Rik | 3573.48735 | 0.47041135 | 4.43E-05 | 0.01682116 |
| Rasgef1b | 745.3761 | 0.46762211 | 0.00013002 | 0.02320938 |
| Smpd3 | 2175.15293 | 0.46364112 | 0.0002248 | 0.0294921 |
| Rragd | 3068.07821 | 0.45981073 | 8.02E-06 | 0.00743786 |
| Tgfa | 1343.72403 | 0.45877183 | 0.00060316 | 0.04311296 |
| Gabra4 | 1964.11491 | 0.45498929 | 5.66E-06 | 0.006678 |
| Filip1 | 543.970132 | 0.45307766 | 0.00029445 | 0.03310177 |
| H2-DMa | 191.461521 | 0.45050294 | 0.00028624 | 0.03310177 |
| Dclk3 | 1754.49477 | 0.45015756 | 0.00022374 | 0.0294921 |
| Diaph2 | 1021.72495 | 0.44075287 | 0.00035263 | 0.03485844 |
| Mctp1 | 2068.58234 | 0.43760904 | 0.00051783 | 0.04204631 |
| Rassf3 | 313.145865 | 0.43535205 | 4.25E-05 | 0.01682116 |
| Epha7 | 756.649629 | 0.42772808 | 0.00051808 | 0.04204631 |
| Pmepa1 | 1018.2751 | 0.42461666 | 0.000487 | 0.04054544 |
| Dgkb | 6501.68119 | 0.42447656 | 0.00054568 | 0.0426947 |
| AI593442 | 2622.75571 | 0.42356477 | 8.84E-05 | 0.01961082 |
| Galnt18 | 346.674992 | 0.42332973 | 0.00067889 | 0.04578551 |
| Kcng1 | 295.350052 | 0.41738485 | 0.00025689 | 0.03149665 |
| Camk2n1 | 10281.142 | 0.41345675 | 6.94E-05 | 0.0195909 |
| Cnksr2 | 2957.18687 | 0.41061142 | 0.00055088 | 0.04271608 |
| Hrk | 679.290714 | 0.40904892 | 0.00014299 | 0.02411898 |
| Pcsk2 | 2957.93364 | 0.40659244 | 3.81E-05 | 0.01681495 |
| 2810459M11Rik | 237.529608 | 0.40496126 | 6.20E-05 | 0.01854398 |
| Shank3 | 4219.96026 | 0.39997231 | 0.00059131 | 0.04311296 |
| Necab2 | 2030.80515 | 0.39982931 | 0.00081774 | 0.04939919 |

| | | | | |
|---|---|---|---|---|
| Dbp | 736.513379 | 0.39535021 | 0.00021885 | 0.0294921 |
| Sema3e | 529.296069 | 0.3940026 | 0.00076994 | 0.04761894 |
| Ppp3ca | 22331.1738 | 0.39296117 | 0.00028576 | 0.03310177 |
| Trnp1 | 1029.98508 | 0.39043348 | 0.00042731 | 0.03801291 |
| Cplx2 | 18164.0609 | 0.38001612 | 0.00011495 | 0.02296941 |
| Mef2c | 2438.45151 | 0.37965201 | 1.82E-05 | 0.01075937 |
| Chn1 | 7318.8937 | 0.37630448 | 0.00073774 | 0.04735542 |
| Dgat2 | 983.796304 | 0.3740808 | 0.00034364 | 0.03482863 |
| Osbp2 | 703.910916 | 0.36823078 | 0.00072197 | 0.04688472 |
| Ppp1r16b | 3338.82068 | 0.36412481 | 0.00079695 | 0.04859495 |
| Arhgap32 | 5624.92538 | 0.36279139 | 0.00023487 | 0.03043355 |
| Gpr158 | 2826.27972 | 0.36139748 | 0.00058668 | 0.04311296 |
| Id3 | 259.698703 | 0.36069307 | 5.74E-05 | 0.01818702 |
| Id4 | 4326.09754 | 0.35930696 | 0.00012264 | 0.02308458 |
| Rapgef4 | 8141.75275 | 0.35502261 | 6.89E-05 | 0.0195909 |
| Smad3 | 1082.66417 | 0.35270375 | 0.00038448 | 0.03657472 |
| Mgat5b | 1019.8715 | 0.35185315 | 0.00055253 | 0.04271608 |
| Prom1 | 246.83504 | 0.35037762 | 1.82E-05 | 0.01075937 |
| Zfp365 | 5459.89321 | 0.34856326 | 0.00034105 | 0.03482863 |
| Spock3 | 3993.33259 | 0.34781457 | 0.0001857 | 0.02857924 |
| Rtn4rl1 | 587.572245 | 0.34588045 | 0.00038727 | 0.03657472 |
| Hlf | 2539.95263 | 0.34476295 | 9.69E-06 | 0.00839322 |
| Slmap | 2776.61714 | 0.34143709 | 0.00031787 | 0.03411993 |
| Calb1 | 4796.27589 | 0.34106082 | 8.08E-05 | 0.01961082 |
| Cdkl5 | 1015.31858 | 0.33460408 | 0.00026609 | 0.03229939 |
| Rhobtb2 | 1725.37313 | 0.33446415 | 0.00025706 | 0.03149665 |
| Phyhip | 6655.11847 | 0.33110168 | 0.00065623 | 0.04524505 |
| Ppp1r9a | 9848.81834 | 0.32928282 | 5.66E-05 | 0.01818702 |
| Arl15 | 593.051519 | 0.32748503 | 0.00047759 | 0.0402792 |
| Ppp1r9b | 10681.3982 | 0.3271117 | 0.00071407 | 0.04683975 |
| Mapk4 | 1311.42041 | 0.32599928 | 0.00017893 | 0.02834039 |
| Ppp4r4 | 988.324998 | 0.32483387 | 0.00082581 | 0.04942682 |
| Wasf1 | 3132.43582 | 0.32276165 | 0.00049792 | 0.04119083 |
| Plppr1 | 743.7781 | 0.32155768 | 0.00066385 | 0.04524505 |
| Agfg2 | 731.644952 | 0.32112413 | 0.00027153 | 0.03235492 |
| Tomm70a | 5526.20344 | 0.31858601 | 0.00015662 | 0.0260791 |
| Ndrg4 | 26119.3655 | 0.31738018 | 0.00066537 | 0.04524505 |
| C2cd2l | 3515.8293 | 0.31534187 | 0.00075855 | 0.04735542 |
| Gucy1b1 | 2847.13449 | 0.30096454 | 0.00013379 | 0.02320938 |
| Ppp1ca | 2425.56621 | 0.30002231 | 0.00027081 | 0.03235492 |
| Igsf11 | 816.756546 | 0.29617933 | 0.00025135 | 0.03138945 |

| | | | | |
|---|---|---|---|---|
| Neurl1a | 2882.99609 | 0.29450672 | 0.00070622 | 0.04683975 |
| Kcnh5 | 287.653198 | 0.29437061 | 0.00044775 | 0.03902923 |
| Tacc1 | 3594.80785 | 0.29335227 | 0.00029564 | 0.03310177 |
| Gas7 | 7614.06439 | 0.29222293 | 0.00029564 | 0.03310177 |
| Prkcz | 3122.70415 | 0.29187016 | 0.0001954 | 0.02859076 |
| Lrrc20 | 323.737075 | 0.28846963 | 0.00053604 | 0.04245156 |
| Tnfrsf19 | 718.207251 | 0.27969728 | 0.00019592 | 0.02859076 |
| Cxcl12 | 784.551971 | 0.27736749 | 4.29E-05 | 0.01682116 |
| Cdk19 | 1662.16024 | 0.27126261 | 5.47E-05 | 0.01818702 |
| Gpr155 | 1548.8982 | 0.26683141 | 0.00035341 | 0.03485844 |
| Adcy1 | 2300.47846 | 0.26192307 | 0.00059297 | 0.04311296 |
| Tmem132b | 2428.61821 | 0.25428606 | 1.51E-05 | 0.01075937 |
| Pcdh10 | 5169.09474 | 0.25346096 | 0.00042453 | 0.03801291 |
| Galnt13 | 1448.01691 | 0.25335311 | 4.02E-05 | 0.01682116 |
| Sesn3 | 1943.52511 | 0.25309072 | 0.00014049 | 0.02400929 |
| Snn | 2612.65395 | 0.25086045 | 7.91E-06 | 0.00743786 |
| Cntn4 | 523.981115 | 0.24537057 | 0.0006093 | 0.04324385 |
| Mkrn1 | 1585.58225 | 0.24401552 | 0.00030488 | 0.03382479 |
| Mras | 2668.27501 | 0.24212837 | 0.00071393 | 0.04683975 |
| Zbtb7a | 2185.75756 | 0.23670726 | 0.00072979 | 0.04715697 |
| Sox9 | 787.671076 | 0.23575877 | 0.00030762 | 0.03382479 |
| Frmpd4 | 1614.95042 | 0.23296982 | 3.76E-05 | 0.01681495 |
| Mtmr12 | 455.789119 | 0.2327033 | 0.00081738 | 0.04939919 |
| Orai2 | 997.659326 | 0.23203775 | 0.00018413 | 0.02857924 |
| Clip4 | 1478.5019 | 0.23162781 | 0.00035427 | 0.03485844 |
| Reps2 | 3696.81793 | 0.22812785 | 0.0007969 | 0.04859495 |
| Ubtf | 1998.26909 | 0.22619851 | 0.00011243 | 0.02281618 |
| Cldn12 | 956.835158 | 0.22151282 | 0.00038861 | 0.03657472 |
| St3gal5 | 1258.30182 | 0.21798829 | 0.00059009 | 0.04311296 |
| Chmp3 | 1627.82401 | 0.2136589 | 0.00046071 | 0.03936811 |
| Tef | 5198.64878 | 0.21334509 | 0.00042344 | 0.03801291 |
| Dennd11 | 2204.05322 | 0.20952521 | 0.00023701 | 0.03043355 |
| Ccny | 2051.1511 | 0.20477054 | 3.65E-05 | 0.01681495 |
| Mxi1 | 1288.37948 | 0.20301146 | 0.00068037 | 0.04578551 |
| Bhlhe41 | 995.433714 | 0.20265658 | 0.00041447 | 0.0376445 |
| Zhx1 | 1909.88704 | 0.19612428 | 0.00053334 | 0.04245156 |
| Snw1 | 1140.23397 | 0.19529783 | 0.00065955 | 0.04524505 |
| Gpam | 1746.8653 | 0.16816227 | 0.00036033 | 0.03511653 |
| Dpysl2 | 2963.42914 | -0.1670872 | 0.00056484 | 0.04284925 |
| Clk3 | 782.75105 | -0.1904031 | 0.00046073 | 0.03936811 |
| Prpsap1 | 644.673504 | -0.1986134 | 0.00060414 | 0.04311296 |

| | | | | |
|---|---|---|---|---|
| Tbkbp1 | 852.688219 | -0.2245414 | 0.00069236 | 0.04635247 |
| Sorcs3 | 1023.94009 | -0.22523 | 0.00013118 | 0.02320938 |
| Uba5 | 813.71026 | -0.2323316 | 8.90E-05 | 0.01961082 |
| Sema4a | 806.713826 | -0.2401795 | 0.00076111 | 0.04735542 |
| Taf15 | 801.940131 | -0.24158 | 0.00029045 | 0.03310177 |
| Polr3e | 663.06544 | -0.2448902 | 8.22E-05 | 0.01961082 |
| Brd9 | 780.420367 | -0.2583753 | 0.00021792 | 0.0294921 |
| Grk3 | 1636.50064 | -0.2649624 | 0.00061967 | 0.04374024 |
| Peg10 | 10281.2001 | -0.2656872 | 0.00075943 | 0.04735542 |
| Epha5 | 1289.14926 | -0.2674514 | 0.00059959 | 0.04311296 |
| Ankrd13d | 760.387991 | -0.2719043 | 0.00056494 | 0.04284925 |
| Arid5b | 613.423799 | -0.2746477 | 0.00075943 | 0.04735542 |
| Arrb2 | 563.845426 | -0.3132115 | 1.58E-05 | 0.01075937 |
| Plk3 | 177.695522 | -0.3286053 | 0.00075024 | 0.04735542 |
| F3 | 692.43367 | -0.3369645 | 7.22E-06 | 0.00743786 |
| Alk | 278.715808 | -0.3421647 | 0.00034593 | 0.03482863 |
| Hdac7 | 528.999561 | -0.3422681 | 0.00071057 | 0.04683975 |
| Prr5 | 249.217694 | -0.3443065 | 6.24E-05 | 0.01854398 |
| Otof | 1138.61979 | -0.3466157 | 0.00056204 | 0.04284925 |
| Slc5a3 | 1408.77153 | -0.3474649 | 3.28E-06 | 0.00607686 |
| Ankrd24 | 566.679767 | -0.357104 | 0.00046418 | 0.03940336 |
| Baiap3 | 6050.13989 | -0.3588317 | 0.00016552 | 0.02653963 |
| Gpr161 | 319.80106 | -0.3673266 | 1.98E-05 | 0.01117013 |
| Ebf4 | 231.53983 | -0.3845785 | 0.00016192 | 0.02628802 |
| Cpne7 | 1531.05236 | -0.4081035 | 0.00013316 | 0.02320938 |
| Nts | 643.860885 | -0.4087929 | 0.00021983 | 0.0294921 |
| Zfp963 | 96.5646275 | -0.4186095 | 0.00050851 | 0.0418009 |
| Vamp1 | 1002.66932 | -0.4242631 | 8.18E-05 | 0.01961082 |
| Tsc22d3 | 1004.29073 | -0.4394077 | 2.47E-05 | 0.01335128 |
| Trib1 | 141.374419 | -0.4445363 | 0.00012017 | 0.02308458 |
| Lsm11 | 208.058195 | -0.4598408 | 1.87E-06 | 0.00501306 |
| Col15a1 | 140.550134 | -0.4631059 | 0.00046057 | 0.03936811 |
| Vwa5b2 | 604.669435 | -0.4726503 | 0.00018809 | 0.02857924 |
| Fabp7 | 339.835545 | -0.480844 | 0.00075045 | 0.04735542 |
| Crh | 280.43959 | -0.4851822 | 0.00052121 | 0.04204631 |
| Tmem145 | 445.731263 | -0.4869063 | 0.00082551 | 0.04942682 |
| Col16a1 | 363.126433 | -0.4952326 | 8.91E-05 | 0.01961082 |
| Ptpru | 536.436584 | -0.4977095 | 4.58E-06 | 0.00661422 |
| Ankrd23 | 88.0230498 | -0.49917 | 0.00022176 | 0.0294921 |
| Cgn | 194.949345 | -0.5088331 | 0.00065517 | 0.04524505 |
| Akap12 | 487.340177 | -0.5109548 | 4.84E-05 | 0.01745418 |

| | | | | |
|---|---|---|---|---|
| Col11a1 | 342.679762 | -0.5305025 | 0.00059984 | 0.04311296 |
| Mapk11 | 180.127401 | -0.5352577 | 0.00012103 | 0.02308458 |
| Adcy7 | 258.040789 | -0.5436081 | 1.41E-05 | 0.01075937 |
| Slc1a6 | 156.466445 | -0.5518564 | 0.00013402 | 0.02320938 |
| Gdpd2 | 280.670279 | -0.6154154 | 7.91E-05 | 0.01961082 |
| Gm13112 | 92.7673967 | -0.6156006 | 2.77E-05 | 0.01437017 |
| Mamdc4 | 56.803838 | -0.616274 | 0.00013121 | 0.02320938 |
| Ttc39aos1 | 108.898025 | -0.6585685 | 0.00021007 | 0.0294921 |
| Coro6 | 264.713423 | -0.6778911 | 0.0001028 | 0.02153442 |
| Zfp57 | 236.193702 | -0.6821946 | 9.43E-07 | 0.00408363 |
| Arrdc2 | 155.914339 | -0.7151582 | 3.17E-10 | 4.12E-06 |
| Cdh23 | 69.6004476 | -0.7812334 | 0.00038389 | 0.03657472 |
| Sgk1 | 1207.86131 | -0.8415739 | 1.93E-06 | 0.00501306 |
